# ATF4 mediates fetal globin upregulation in response to reduced β-globin

**DOI:** 10.1101/2020.01.15.905943

**Authors:** Mandy Boontanrart, Gautier Stehli, Marija Banovic, Markus S. Schröder, Stacia Wyman, Rachel Lew, Matteo Bordi, Benjamin Gowen, Mark DeWitt, Jacob E. Corn

**Affiliations:** Department of Molecular and Cell Biology, University of California Berkeley. Berkeley, CA, USA; Department of Biology, ETH Zurich. Zurich, Switzerland; Innovative Genomics Institute, University of California Berkeley. Berkeley, CA, USA; Spotlight Therapeutics. Hayward, CA, USA

## Abstract

Fetal development and anemias such as β-hemoglobinopathies trigger rapid production of red blood cells in a process known as stress erythropoiesis. Cellular stress prompts differentiating erythroid precursors to express high levels of fetal γ-globin, which has suggested strategies to treat hemoglobinopathies such as thalassemia and sickle cell disease. However, the mechanisms underlying γ-globin production during cellular stress are still poorly defined. Here we use CRISPR-Cas genome editing and CRISPRi transcriptional repression to model the stress caused by reduced levels of adult β-globin. We find that loss of β-globin is sufficient to induce widespread globin compensation, including robust re-expression of γ-globin. Time-course RNA-seq of differentiating isogenic erythroid precursors identified the ATF4 transcription factor as a causal regulator of this response. ChIP-seq of multiple erythroid precursor genotypes and differentiation states revealed that β-globin knockout leads to reduced engagement of ATF4 targets involved in the unfolded protein response. This ATF4 program indirectly regulates the levels of BCL11A, a key repressor of γ-globin. Identification of ATF4 as a key regulator of globin compensation adds mechanistic insight to the poorly understood phenomenon of stress-induced globin compensation and could be relevant for proposed gene editing strategies to treat hemoglobinopathies.

## Introduction

Red blood cells (RBCs), also known as erythrocytes, are packed with hemoglobin tetramers and circulate throughout the body to supply all tissues with oxygen. Adult RBCs primarily contain adult hemoglobin (HbA), which consists of two copies of α-globin and two copies of β-globin. β-thalassemic genetic disorders are caused by disruption of β-globin expression, causing loss of HbA and resulting in severe anemia, poor growth, and dramatically shortened lifespan. β-thalassemia can be ameliorated by re-expression of γ-globin, which complexes with α-globin to form fetal hemoglobin (HbF). γ-globin is normally expressed during development and silenced soon after birth, in an inverse relationship with β-globin.

Severe anemias such as β-thalassemia can trigger rapid production of red blood cells in a process known as stress erythropoiesis. Stress erythropoiesis is induced by tissue hypoxia resulting from anemia and involves a distinct erythropoietic program that favors increased γ-globin. For example, recovery from bone marrow transplant or other treatments that greatly reduce erythroblast levels is often characterized by high levels of γ-globin and HbF (Alter, 1979; Meletis et al., 1994; Papayannopoulou et al., 1980; Weinberg et al., 1986). Homozygous β-thalassemia patients transplanted with bone marrow from heterozygous siblings express high levels of HbF that can be sustained for up to two years after transplant (Galanello et al., 1989). β-thalassemia patients can also re-express high levels of γ-globin without transplant (Manca and Masala, 2008; Rochette et al., 1994). Intrinsic cellular processes can also mimic stress erythropoiesis, and ex *vivo* cultured stress erythroid progenitors express high levels of HbF (Xiang et al., 2015). Overall, this has led to the suggestion that erythroid stress recapitulates fetal erythropoiesis, but the pathways involved in fetal globin expression during this process remain to be determined.

Cell-intrinsic mimicry of fetal erythropoiesis is being actively explored as a potential route to treat a variety of globinopathies, including β-thalassemia and sickle cell disease (Platt et al., 1984; Wienert et al., 2018). These approaches stem from the observation that naturally occurring mutations which promote HbF re-expression can ameliorate β-globinopathy symptoms (Berry et al., 1992; Jacob and Raper, 1958). A variety of strategies for HbF re-expression are being pursued, including large deletions within the β-globin locus, mutations of the *HBG1/2* promoter, and cell-specific reduction of the BCL11A repressor by enhancer editing (Sankaran, 2011; Wienert et al., 2018).

Here, we sought to model the cellular erythroid stress occurring after disruption of β-globin, hereafter referred to as β_0_-stress, in order to determine the mechanism underlying spontaneous re-expression of fetal hemoglobin. We previously found that fetal hemoglobin was upregulated after genome editing of *HBB* in CD34+ adult mobilized hematopoietic stem and progenitor cells (HSPCs) to reverse the causative allele of sickle cell disease. *In vitro* differentiation of bulk edited cell populations induced high *HBG1/2* transcript levels as compared to unedited cells, which translated to high levels of HbF tetramers (DeWitt et al., 2016). This effect persisted even after long-term xenotransplantation of edited cells to immunodeficient mice (https://www.biorxiv.org/content/10.1101/432716v6).

We now show that β_0_-stress caused by reductions in β-globin using either CRISPR-Cas genome editing or CRISPRi transcriptional repression are sufficient to induce high levels of γ-globin in immortalized hematopoietic progenitors and adult mobilized CD34+ HSPCs. Time-course transcriptomics of isogenic *HBB* knockout and wild type cells during erythroid differentiation reveal that loss of β-globin leads to a transcriptional program that induces very high levels of *HBG* and moderately activates the transcription of other globins such as *HBE* and *HBZ.* Pathway analysis indicates that β-globin knockout cells reduce *ATF4* (activating transcription factor 4) activity, leading to reductions in transcripts of many *ATF4* targets. Endogenous mutation of *ATF4* leads to upregulation of several globins, especially γ-globin, much like HBB knockout or knockdown. BCL11A levels were reduced in cells with knockout of *HBB* or an *ATF4* mutation, leading us to first suspect *BCL11A* as a possible ATF4 target.

However, ChIP-seq showed no evidence for ATF4 binding anywhere near *BCL11A* regardless of differentiation status or *HBB* genotype. Instead, we find evidence for involvement of ATF4 in a CEBP coordination program in the undifferentiated state, which switches to a distinct GATA program during differentiation. *HBB* knockout reduces ATF4 binding at multiple genes involved in the unfolded protein response, leading to their transcriptional repression. Overall, our data indicate that β_0_-stress inhibits an ATF4-mediated transcriptional program. Reduction of ATF4 activity indirectly lowers *BCL11A* expression, prevents overall globin repression by the unfolded protein response, thereby upregulating multiple globins especially γ-globin. These data provide mechanistic insight into the long-observed but poorly-understood phenomenon of fetal globin expression during cell-intrinsic erythroid stress.

## Results

### Loss of *HBB* leads to upregulation of γ-globin

We previously observed that CRISPR-Cas9 editing at *HBB* in CD34+ mobilized peripheral blood HSPCs induced γ-globin transcription relative to unedited cells, leading to the formation of HbF tetramers (DeWitt et al., 2016). To mechanistically investigate how β_0_-stress caused by knockout of *HBB* upregulates *HBG,* we used the HUDEP-2 cord-blood derived erythroid progenitor cell line (Kurita et al., 2013). HUDEP-2 cells normally express high levels of *HBB* and low levels of *HBG,* making them a popular model for the study of adult globin regulation (Bauer et al., 2013; Grevet et al., 2018; Wienert et al., 2015).

We made a clonal HUDEP-2 line with homozygous knockout of *HBB* (HBBko) using electroporation of a CRISPR-Cas9 RNP complexed with a well-validated HBB-targeting guide “e66”, which targets the exonic (e) region 66 base pairs from the start of *HBB* (DeWitt et al., 2016) (**Fig S1A**). HBBko was derived from a wild type (WT) HUDEP-2 clone (previously published as H2.1) in order to control for clonal effects associated with the heterogeneous HUDEP-2 parental population (Chung et al., 2019; Wienert et al., 2015). γ-globin levels were strikingly increased in the HBBko line by intracellular flow cytometry, with the vast majority of edited cells expressing detectable levels of the protein (**Fig 1A**).

**Figure 1.**
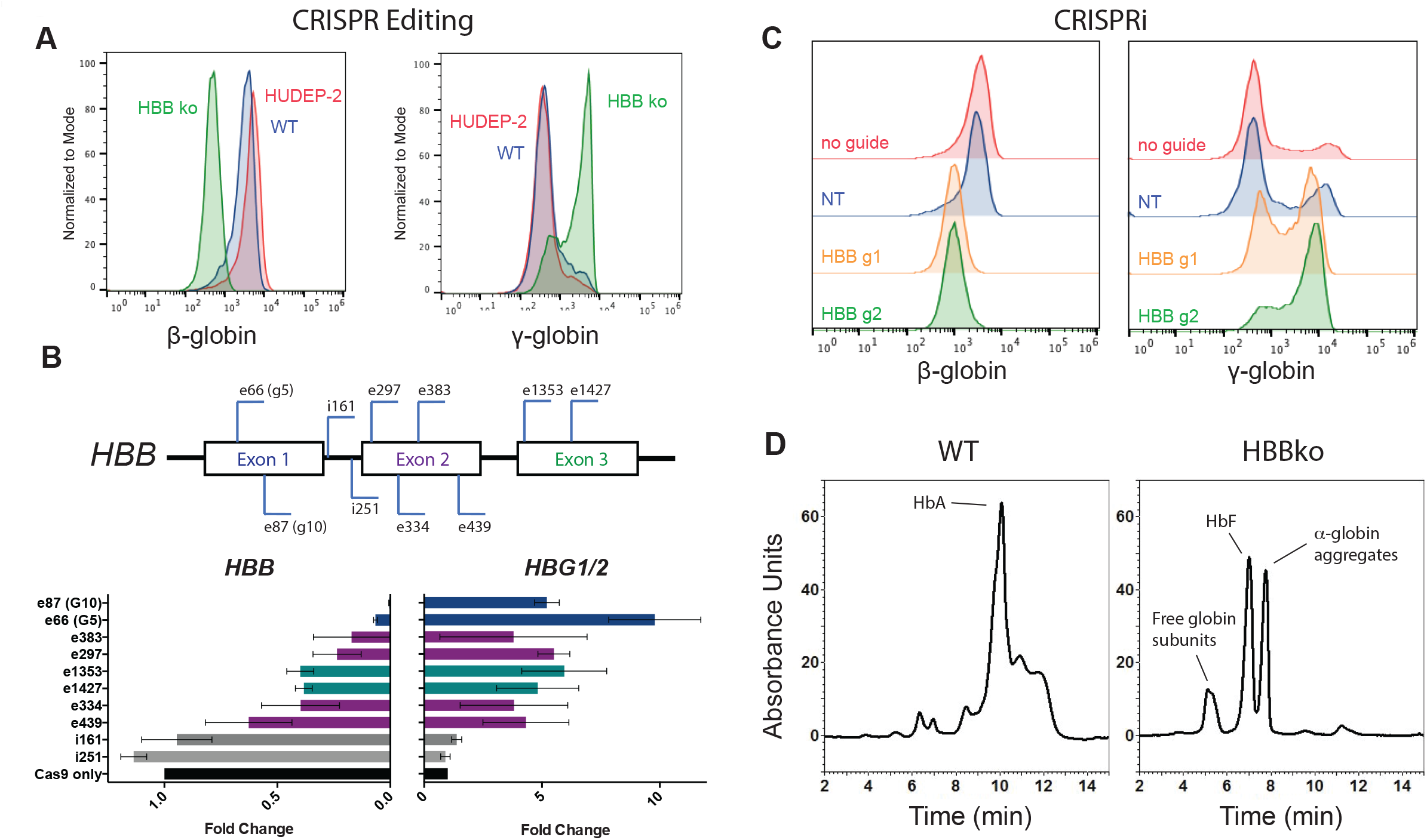
Loss of β-globin leads to increased γ-globin in HUDEP-2 cells and CD34+ HSPCs. A) A *HBB* knockout (HBBko) line was generated from a wild type (WT) HUDEP-2 subclone though Cas9 ribonucleoprotein (RNP) targeting of exon 1 of *HBB.* Intracellular flow cytometry staining of β-globin and γ-globin of differentiated HUDEP-2 pool, WT clone, and HBBko clone. B) Schematic of sgRNAs targeting *HBB* at either exonic (e) or intronic (i) positions. Numbers indicate the number of bases each RNP would cut after the transcription start site of the *HBB* gene. Previously published “G5” and “G10” sgRNAs are noted (DeWitt et al., 2016). Below is qRT-PCR of *HBB* and *HBG1/2* after pooled editing of the WT line with each sgRNA and 5 days of differentiation. Editing efficiency of each sgRNA is shown in **Fig S1D**. The data is presented as mean ± SD of three biological replicates. C) WT HUDEP-2 cells stably expressing dCas9-KRAB with guides targeting HBB were differentiated for 5 days and hemoglobin levels were measured by intracellular flow cytometry. D) WT and HBBko cells were differentiated for 5 days and hemolysates were analyzed by HPLC. Peak compositions were identified through mass spectrometry (**Table S1**).

Both the WT and HBBko lines were equally well *in vitro* differentiated to erythroblasts under standard conditions, as measured by staining for CD235a (Glycophorin A). WT and HBBko cells exhibited no apparent difference in cell survival during bulk differentiation, suggesting that β_0_-stress induces γ-globin expression in the majority of cells as opposed to conferring a fitness advantage towards cells that stochastically express more γ-globin (**Fig S1B**). WT expressed high levels of β-globin after differentiation, but as expected HBBko completely lost β-globin protein by intracellular FACS. *HBB* mRNA was still detectable by qRT-PCR in HBBko, but was slightly reduced relative to WT in undifferentiated cells and 95% reduced in differentiated cells (**Fig S1C**).

DNA damage associated with CRISPR-Cas9 genome editing has been linked to long-lasting cellular phenotypes in HSPCs, including p53 activation (Schiroli et al., 2019). To distinguish non-specific effects of genome editing from specific intervention at the β-globin locus, we tested a panel of Cas9 RNPs complexed with *HBB* guide RNAs targeting exonic or neighboring intronic regions of *HBB* (**Fig. 1B**). After bulk editing and *in vitro* differentiating the WT line, we found that multiple RNPs targeting coding regions consistently reduced *HBB* levels and increased *HBG,* as measured by qRT-PCR (**Fig 1B**). RNPs targeting neighboring intronic regions neither decreased *HBB* nor increased *HBG* despite similar levels of editing (**Fig S1D**). We found similar results when editing and *in vitro* differentiating CD34+ adult mobilized HSPCs from multiple donors with a panel of exonic or intronic HBB-targeting Cas9 RNPs (**Fig S1E** and **Fig S1F**). Bulk-editing *HBB* in CD34+ HSPC cells did not significantly alter cell numbers during expansion (**Fig S1G**). Additionally, total cell numbers after *in vitro* differentiation were comparable amongst exonic and intronic HBB-targeting guides (**Fig S1H**). As with isogenically edited HUDEP-2 cells, this implies that β_0_-stress in HSPCs induces γ-globin expression rather than positively selecting for cells that already express γ-globin.

The ability of multiple HBB-targeting RNPs to induce *HBG* expression suggested that this effect was independent of the genomic change and was instead a response to loss of *HBB* transcript. To test this hypothesis, we made a stable WT HUDEP-2 subclone expressing dCas9-KRAB (CRISPRi) (**Fig S1I**). We found that stable CRISPRi of *HBB* using two different guide RNAs led to potent downregulation of β-globin and upregulation of γ-globin protein by intracellular flow cytometry (**Fig 1C**). This was also reflected by levels of *HBB* and *HBG* transcripts as measured by qRT-PCR (**Fig S1J**). Overall, these data indicate that upregulation of *HBG* is not a consequence of DNA damage or genomic targeting, but is induced by loss of *HBB*.

To better characterize the globin protein composition resulting from loss of *HBB,* we used high performance liquid chromatography (HPLC) coupled to mass spectrometry (MS) of *in vitro* differentiated HUDEP-2 clones. We separated intact globin peaks using standard HPLC conditions while also collecting fractions. We subjected each fraction to mass spectrometry in order to unambiguously assign the globin composition of each peak. HPLC of the WT clone resulted in a peak pattern suggesting a typically “adult” set of globins, including high levels of HbA (**Fig 1D**. This peak assignment was confirmed on the molecular level by mass spectrometry (**Table S1**). HPLC of the HBBko clone instead revealed a very different pattern of peaks (**Fig 1D**). Mass spectrometry and comparison to literature globin HPLC led us to assign these as uncomplexed globins, α-globin aggregates, and HbF tetramers (Lechauve et al., 2019) (**Table S1**).

Our data thus far indicate that low levels of *HBB* caused by either genome editing or CRISPRi are associated with high levels of *HBG.* The presence of α-globin aggregates in cells with β_0_-stress suggests that an imbalance of hemoglobin proteins might lead to an unfolded protein response.

We therefore asked whether the relationship between levels of *HBB* and *HBG* was specific for reductions in *HBB* or a reflected an issue of general globin balance. For this, we utilized HUDEP-1 cells, which are similar to HUDEP-2s but exhibit a mostly immature globin profile of low *HBB* and high *HBG.* Pooled knockout of *HBG* using a CRISPR-Cas RNP led to loss of *HBG* mRNA and upregulation of *HBB* (**Fig S1K**). Therefore, our data suggest that erythroid progenitors sense globin levels during differentiation and attempt to compensate for missing globins by upregulating what they can. This model is consistent with the phenomenon of increased γ-globin during cell-intrinsic erythroid stress, and indicates that β_0_-stress modeled by the HBBko line is useful to molecularly characterize this process.

### HBB knockout cells exhibit ATF4-mediated transcriptional reprogramming during differentiation

Because *HBB* and *HBG* reciprocally regulated one another, we asked if *HBB* loss affects the transcription of other globins. We performed biological triplicate *in vitro* differentiation of paired WT and HBBko clones. RNA-seq of terminally differentiated cells revealed widespread transcriptional alterations between WT and HBBko, with some of the largest fold changes being greatly reduced *HBB* and increased *HBG1, HBG2, HBE, and HBZ* (**Fig 2A**). There were relatively low total levels of *HBE* and *HBZ* in even HBBko cells after differentiation, such that *HBG1/2* comprised the vast majority of globin transcripts.

**Figure 2.**
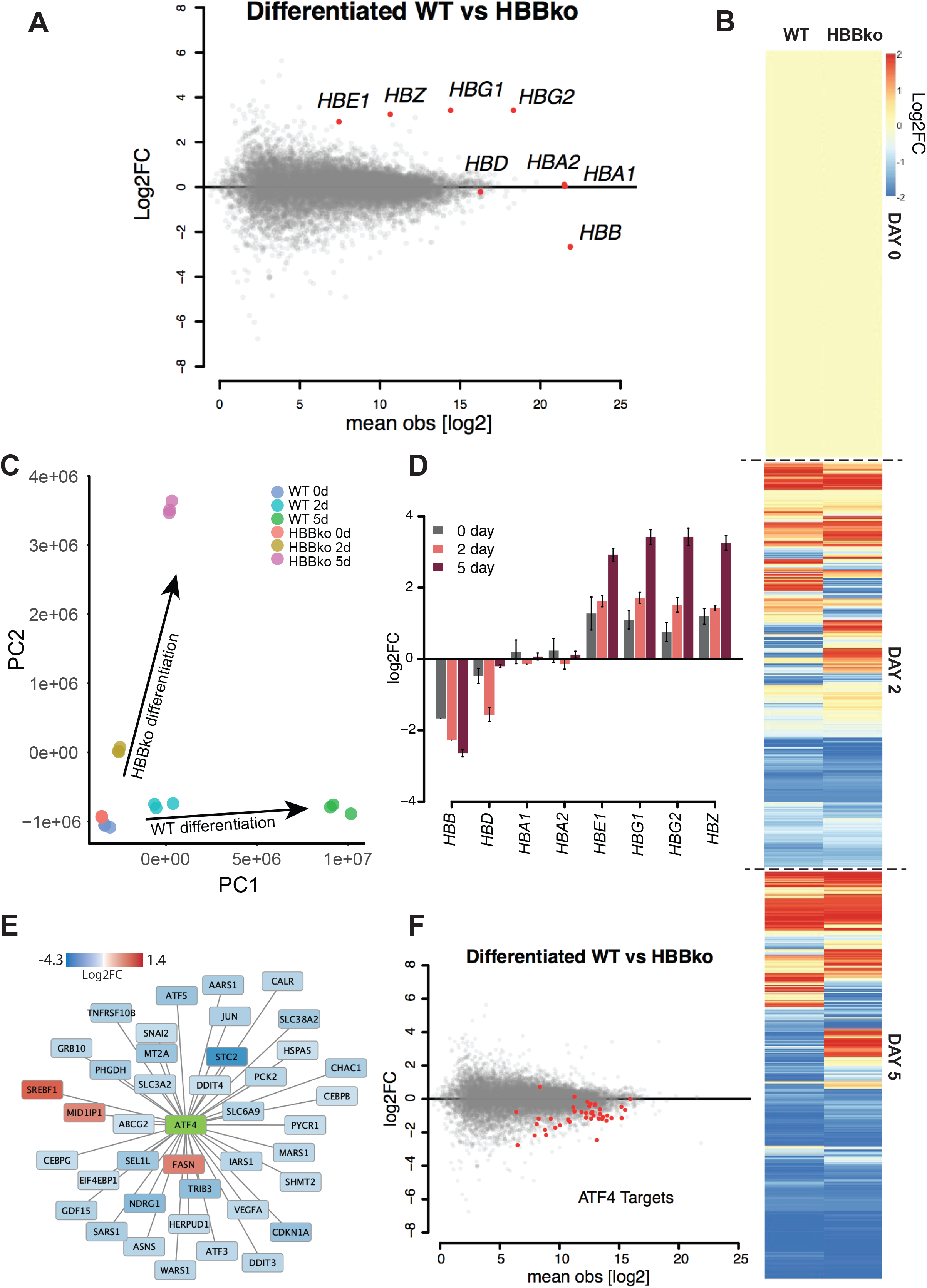
Transcriptomics of time-course differentiation reveals an ATF4 signature in HBBko cells. A) MA plot after 5 days of differentiation comparing HBBko and WT cells. Globin genes are highlighted in red. RNA-seq was performed in biological triplicates and data is shown as the mean. B) Biological triplicate RNA-seq data of WT and HBBko cells undifferentiated (day 0), 2 days of differentiation, and 5 days of differentiation. Transcript levels are expressed as log2 fold-change normalized to their respective day 0 expression. C) Principal component analysis of all RNA-seq samples shows agreement of replicates and alterations in differentiation trajectories between WT and HBBko. Replicated colors indicate n=3 biological replicates of each condition. D) The expressions of embryonic, fetal, and adult globin genes are plotted for the 3 timepoints of differentiation. Data is shown as log2 fold change relative to WT cells and presented as mean ± SD of 3 biological replicates. E) Pathway analysis identifies 38 out of 40 ATF4 targets as significantly affected in HBBko cells. Genes highlighted in blue and red are ATF4 targets that are downregulated and upregulated genes by log2 fold change on Day 5 of differentiation, respectively. F) MA plot from 5 days of differentiation highlighting 40 known ATF4 targets (red). The mean of biological triplicates is shown.

To capture the dynamics of differentiation, we performed triplicate time course RNA-seq by collecting samples in their undifferentiated state, after two days of differentiation, and after five days of differentiation (**Fig S2A-B**). Globally, the transcriptomes of WT and HBB knockout cells are similar throughout differentiation (**Fig S2C**). However, on top of the overall similarity, the knockout of *HBB* induces a distinct transcriptional program that emerged after two days of differentiation and intensified by five days (**Fig 2B**). This was reflected in PCA analysis, showing that WT and HBBko cells start quite similar, but have distinct differentiation trajectories (**Fig 2C**). Specifically examining the expression of globin genes, we found that as *HBB* expression declines during differentiation in HBBko cells as compared to WT, other globins become more highly expressed (**Fig 2D**).

Nonsense-Induced Transcriptional Compensation (NITC) is a response to nonsense-mediated decay (NMD) of an important transcript and has emerged as a new mechanism for transcriptional reprogramming to upregulate related genes (El-Brolosy et al., 2019; Ma et al., 2019; Rossi et al., 2015). However, we found that knocking down the mRNA decay exonuclease XRN1 or the single-stranded RNA binding protein SMG6 that are required for NITC had no effect on the upregulation of γ-globin in HBBko cells by intracellular FACS and qRT-PCR (**Fig S2D**).

We used pathway analysis (Krämer et al., 2014) to identify features in the RNA-seq data that could explain the transcriptional reprogramming in differentiated HBBko cells. We found that ATF4 (Activating Transcriptional Factor 4) and 37 out of 40 ATF4 target genes were downregulated in HBBko cells relative to WT cells (**Fig 2E, F**).

ATF4 is a bZIP family transcription factor associated with the integrated stress response (ISR) to unfolded proteins. ATF4 signaling decreases global protein synthesis while simultaneously inducing target genes. The downregulation of ATF4 and its ISR targets in HBBko cells was therefore surprising since these cells harbor free α-chains that might have conversely initiated the ISR (**Fig 1D**) (Chen and Zhang, 2019). However, a focused CRISPR screen recently found that knockout of the known ATF4 upstream regulator HRI (heme regulated eIF2a kinase, *EIF2AK1)* can induce moderate levels of HbF expression (Grevet et al., 2018). We hypothesized that both *HBB* and *HRI* knockout may induce globin synthesis by reducing ATF4 activity. This would mark ATF4 as a master regulator whose level maintains globin levels at homeostasis when a major globin becomes critically low.

### ATF4 represses fetal hemoglobin expression

We developed two CRISPR-Cas9 genome editing strategies to test whether ATF4 was involved in globin homeostasis, and specifically in fetal globin regulation in cells that predominantly express adult globin. First, we used a dual-guide approach to excise the entirety of the *ATF4* gene, starting from the WT HUDEP-2 clone (**Fig S3A**). Two ATF4 knockout clones resulting from total excision were named ATF4ko-1 and ATF4ko-2 (**Fig S2B**). The knockout clones showed successful knockout of ATF4 by Western blotting, with no protein detected even after treatment with cyclopiazonic acid (CPA), a drug that increases ATF4 expression through induction of endoplasmic reticulum stress (**Fig S3C**). Notably, ATF4ko-1 and ATF4ko-2 were unable to undergo *in vitro* differentiation, and instead exhibited extensive cell death after 5 days in differentiation media (**Fig S3D**). This is consistent with a known role of *ATF4* in erythroid differentiation (Masuoka and Townes, 2002; Suragani et al., 2012; Zhang et al., 2019).

Second, we explored N-terminal truncation mutants of ATF4 with separation-of-function properties (Steinmüller and Thiel, 2005). We found that dual guides targeted to remove most of the first exon of *ATF4* (**Fig S3A**) lead to reinitiation of translation at the next ATG in the second exon. This produces a stable protein fragment of ~30 kDa lacking the N-terminal regulatory region but retaining the C-terminal DNA binding domain (**Fig S3C**). We derived an edited clone from WT HUDEP-2s using the above strategy, naming it ATF4ΔN (**Fig S3E**). By ChIP-qPCR, we found that ATF4ΔN still binds the *ASNS* promoter but qRT-PCR indicated that ATF4ΔN does not support transcription of *ASNS* (**Fig S3F**). By contrast to ATF4ko-1 and ATF4ko-2, the ATF4ΔN clone was still able to successfully differentiate as measured by high live cell numbers and successful globin expression after 5 days of culture in differentiation media (**Fig S3D, Fig S3G**).

We tested the globin status of the ATF4ko and ATF4ΔN cell lines. Undifferentiated ATF4ko and ATF4ΔN cells expressed normal levels of β-globin and only slightly increased γ-globin by intracellular FACS (**Fig 3A, Fig S3H**) and Western Blot (**Fig 3B**). Strikingly, *in vitro* differentiation of ATF4ΔN led to high levels of transcripts for *HBB, HBE, HBZ,* and *HBG* (**Fig S3I**). This was reflected in increased γ-globin by intracellular FACs (**Fig 3C, Fig S3G**). The global upregulation of globin expression with especially high levels of γ-globin upon differentiation of ATF4ΔN mirrors the response in differentiated *HBB* knockout cells.

**Figure 3.**
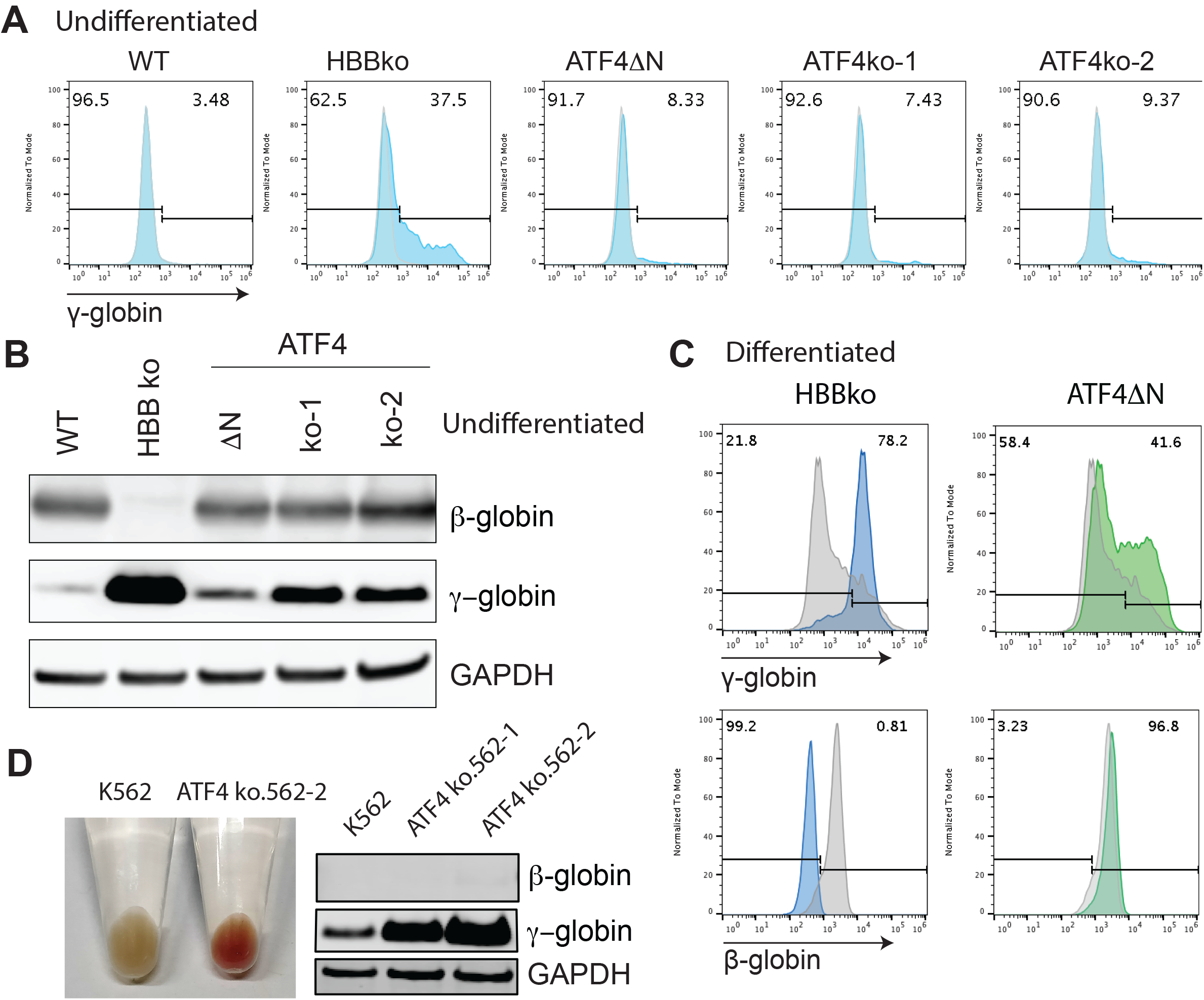
Endogenous mutation of ATF4 leads to increased γ-globin in HUDEP-2 and K562 cells. A) Intracellular FACS for γ-globin of undifferentiated HUDEP2 cells comparing WT, HBBko, and ATF4 clones. Biological quadruplicates are quantified and shown in **Fig S3H**. B) Western blot of undifferentiated WT, HBBko, and ATF4 mutant cells showing β-globin and γ-globin levels. C) Intracellular FACS for γ-globin of differentiated HBBko and ATF4ΔN cells as compared to WT (trace in gray). Biological triplicates are quantified in **Fig S3G**. D) Left panel: cell pellets from wild type K562 and ATF4ko K562 cells. Right panel: Western blot for β-globin and γ-globin in WT and ATF4ko K562 clones. qRT-PCR confirmed loss of *ATF4* transcript in ATF4ko clones and increased *HBG1/2* (**Fig S3J**).

The ability of ATF4 knockout or ATF4ΔN clones to modestly upregulate HbF in the undifferentiated state prompted us to ask whether ATF4 is involved in basal HbF repression in other cell types. We used CRISPR-Cas9 to remove the entire *ATF4* gene from K562 erythroleukemia cells, which normally express no β-globin and low levels of γ-globin. We generated two different ATF4 knockout K562 cell lines that we termed ATF4ko.562-1 and ATF4ko.562-2 using an analogous protocol as for the HUDEP-2 *ATF4* knockout lines (**Fig S3J**). We found that both K562 ATF4 knockout lines became strikingly red and expressed very high levels of γ-globin but still failed to express β-globin (**Fig 3D**), even when they were maintained in undifferentiated culture conditions. Taken together, these data indicate that *ATF4* is involved in repressing fetal hemoglobin in the undifferentiated state, and a lack of *ATF4* signaling in the differentiated state leads to upregulation of multiple globins that mimics the response to lack of *HBB*.

### ATF4 indirectly regulates BCL11A and γ-globin

As a transcription factor, *ATF4* could directly or indirectly regulate the expression of *HBG1/2* and other globins. *BCL11A* and *ZBTB7A* (also known as LRF) are two transcriptional repressors that keep fetal globin levels low after birth. We asked whether BCL11A and ZBTB7A are misregulated in our *HBB* knockout model of β_0_-stress or after loss of *ATF4.*

Examining the WT and HBBko time course differentiation RNA-seq data, we found that ATF4 transcripts were much lower in HBBko in the undifferentiated state, but stabilized to WT levels after 5 days of differentiation. The expression of both *BCL11A* and *ZBTB7A* were reduced in the HBBko line, with *BCL11A* expression declining during differentiation and ZBTB7A remaining at a consistent reduced level (**Fig S4A**). This corresponded to reduced expression of BCL11A protein in HBBko during differentiation (**Fig 4A**). Stable CRISPRi knockdown of *HBB* or *BCL11A* resulted in similar increases in γ-globin as measured by FACS (**Fig S4B**). Knockout or knockdown of HBB both led to a decrease in BCL11A by Western Blot (**Figure 4B**). In the undifferentiated state, the ATF4ΔN and ATF4ko-1 and 2 expressed normal levels of BCL11A and ZBTB7A (**Fig S4C**). However, after differentiation the ATF4ΔN clone exhibited lower levels of BCL11A protein (**Fig 4C**). *HBB* knockout and reduced ATF4 signaling might therefore increase γ-globin expression by directly or indirectly reducing levels of BCL11A during differentiation. However, this is at odds with data from K562 cells, which do not express BCL11A (**Fig S4D**), but dramatically increase *HBG* upon knockout of *ATF4* (**Fig 3D**).

**Figure 4.**
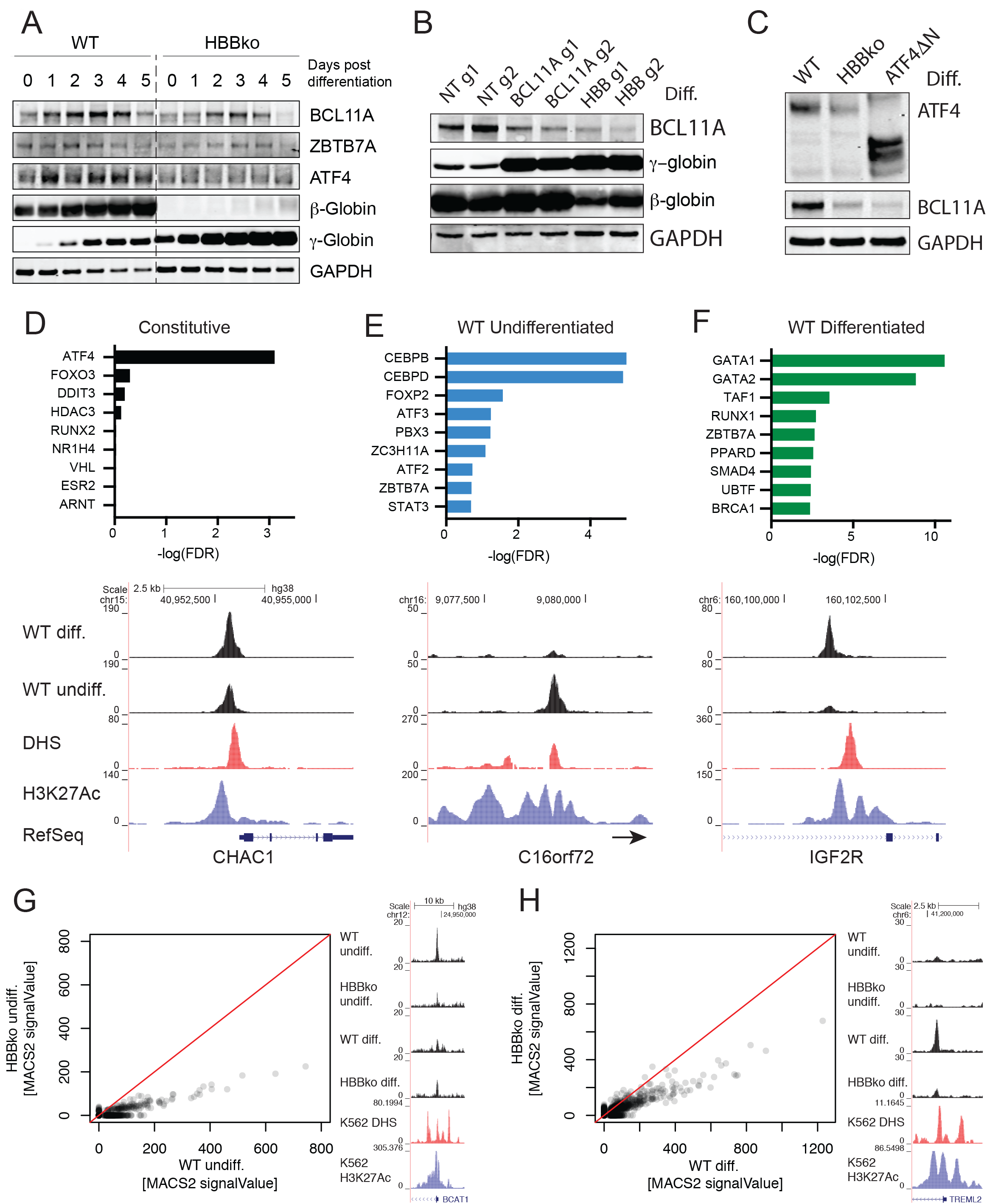
ATF4 indirectly regulates BCL11A and γ-globin. A) Western blot for BCL11A, ZBTB7A, ATF4, β-globin, and γ-globin in HBBko and WT cells over the course of differentiation. B) Western blot showing CRISPRi knockdown efficiency of BCL11A and β-globin, as well as resulting increases in γ-globin protein. C) Western blot of 5 day differentiated WT, HBBko, and ATF4ΔN comparing BCL11A levels. D) Unbiased enrichment analysis of ATF4 ChIP-seq peaks present in both the undifferentiated and differentiated states against databases of transcription factor targets. Example ATF4 ChIP-seq profiles are shown below, together with reference ENCODE DnaseI hypersensitivity and H3K27Ac data. Lack of ATF4 signal at the globin, *BCL11A,* and *ZBTB7A* loci is shown in **Fig S5A-C**. E) Unbiased enrichment analysis of ATF4 ChIP-seq peaks present in only the undifferentiated state. F) Unbiased enrichment analysis of ATF4 ChIP-seq peaks present in only the differentiated state. G) MACS2 signal value for all ATF4 CHIP-seq peaks comparing WT to HBBko cells in the undifferentiated state. Example ATF4 ChIP-seq profiles are shown to the right of each global ChIP-seq comparison. H) MACS2 signal value for all ATF4 CHIP-seq peaks comparing WT to HBBko cells in the differentiated state.

To test whether ATF4 is directly involved in transcriptional regulation of *BCL11A,* we performed endogenous ATF4 ChIP-seq in various HUDEP-2 genetic backgrounds and differentiation states to map ATF4 targets during hemoglobinization. In the undifferentiated state, we performed ATF4 ChIP-seq in WT, HBBko, and ATF4ko (negative control) clones. After five days of *in vitro* differentiation, we performed ATF4 ChIP-seq in WT and HBBko clones, and with an IgG isotype negative control in the WT clone (since the ATF4ko clone cannot be differentiated).

Between all four experimental conditions (WT undifferentiated, HBBko undifferentiated, WT differentiated, HBBko differentiated), we found 1411 unique ATF4 ChIP-seq peaks with an Irreproducible Discovery Rate (IDR) > 2% over background (ATF4 knockout or IgG control).

*Bona fide* ATF4 binding sites showed high read count and clear enrichment over background (e.g up to 255 reads and 85 fold enrichment at *ASNS).* Cross-comparison with our time-course RNA-seq data revealed that ATF4 binding was associated with a subset of well-expressed genes, but ATF4 was not present at unexpressed genes. This further indicates that the identified ATF4 sites are involved in active transcription (**Fig S4E**).

We were surprised to find no evidence of ATF4 binding anywhere in the vicinity of the globin locus, *BCL11A,* or *ZBTB7A* in any sample, including all known promoters and enhancers (**Fig S5A-C**). We therefore used all four ChIP-seq datasets to identify new ATF4 binding sites that varied depending on differentiation status and *HBB* genotype. Unbiased enrichment analysis against databases of transcription factor targets or transcription factor ChIP-seq (Huang et al., 2009; Kuleshov et al., 2016) revealed clear patterns for several distinct categories of ATF4 ChIP-seq peaks.

“Constitutive” ATF4 sites were defined as present regardless of differentiation status in both WT and HBBko. Constitutive peaks were dominated by an ATF4 signal, validating our set of ChIP-seq experiments against datasets in other cell contexts (**Figure 4D, Fig S6A**). “WT Undifferentiated” ATF4 sites were defined as at least two-fold enriched in WT cells in the undifferentiated state and not found in the differentiated state. WT Undifferentiated peaks were highly represented at targets and binding sites of ATF4 partners that either form bZIP heterodimers with ATF4 (e.g. CEBPs) or are part of the same family as ATF4 (e.g. ATF3) (**Figure 4E, Fig S6B**). “WT Differentiated” ATF4 sites were defined as at least two-fold enriched in WT cells in the differentiated state and not found in the undifferentiated state. WT Differentiated peaks were unexpectedly represented at targets and binding sites of transcription factors involved in hematopoietic differentiation (e.g. GATA1/2, and ZBTB7A) but with no known ATF4 partnership (**Figure 4F, Fig S6C**). This latter category implies that ATF4 cooperates with hematopoietic differentiation transcription factors at their target genes, providing a rationale for ATF4’s requirement during hematopoietic differentiation.

Overall, CHIP-seq reads at ATF4 binding sites were attenuated in HBBko cells relative to WT in both the differentiated and undifferentiated states (**Figure 4G-H**). This is consistent with the reduced ATF4 signaling in our timecourse RNA-seq analysis. There are 182 ATF4 sites present in the undifferentiated state that are between two-fold reduced to almost absent in HBBko cells (**Figure 4G**). Only 13 ATF4 sites are present in the differentiated state and are between two-fold reduced or almost absent in HBBko cells (**Figure 4H**). Binding of ATF4 was reduced at several genes involved in the unfolded protein response and feedback regulators of ATF4, including XBP1, HSPA5/BiP, CREB3L1, EDEM1, ATF3, ATF6, RACK1, SEL1L, and TRIB3. Many of these factors are not canonical ATF4 targets, but their expression levels were also reduced in RNA-seq of differentiating HBBko cells (**Fig S7A**). All of these factors are normally involved in reducing protein synthesis during ER stress, and their reduced transcriptional activation by ATF4 could be involved in globin upregulation in HBBko and ATF4ko cells.

## Discussion

Stress erythropoiesis results from a high erythropoietic drive and is induced during anemias such as β-thalassemia. Cell-intrinsic erythroid stress leads upregulation of γ-globin, but the pathways that mediate this phenomenon have remained unclear. We found that β_0_-stress caused by loss of β-globin in erythroid precursors induces ATF4-mediated reprogramming that includes upregulation of several globins including very high levels of γ-globin. ATF4 signaling is required to maintain levels of BCL11A during differentiation, but ATF4 surprisingly does not bind BCL11A or ZBTB7A, nor the globin locus. Hence, we favor a model in which reduced ATF4 signaling during β_0_-stress regulates overall globin levels through an indirect program that may be linked to an attenuated unfolded protein response.

ATF4 is regulated by eIF2-α, which is itself activated by several different kinases including HRI *(EIF2AK1).* Knockout of *EIF2AK1* was recently reported to decrease expression of BCL11A and increase expression of *HBG* (Grevet et al., 2018). However, the link between HRI and BCL11A remained undetermined. Our data suggest that a lack of HRI leads to decreased eIF2-α-phosphorylation and reduction of ATF4 activity, similar to what we observe for knockout of *HBB.* Deletions of *EIF2AK1, HBB,* and *ATF4* all lead to reduced *BCL11A.* Interestingly, deletion of *EIF2AK1* primarily resulted in more *HBG1/2* without substantial changes in other globins or ATF4 targets. By contrast, we find that knockout of *HBB* or *ATF4* leads to upregulation of multiple globins, with *HBG1/2* exhibiting the largest change, as well as reductions in almost all known ATF4 targets. We find that *HBB* knockout leads to reduced ATF4 occupancy at several genes responsible for reduced protein synthesis downstream of ER stress, as well as reduced transcription of each of these targets. Overall, this suggests that ATF4 signaling in response to loss of β-globin is part of a larger program to maintain high globin levels in differentiating erythroid cells. The response to loss of β-globin could include HRI-like signaling as well as additional facets. Knockout of *EIF2AK1* has been suggested as a strategy to ameliorate anemic disorders via re-expression of HbF (Grevet et al., 2018). Indeed, knockout or CRISPRi of *HBB* leads cells to express remarkable levels of *HBG1/2.* But the broad roles of ATF4 in processes other than erythropoiesis also suggest that this strategy should be pursued with caution.

The HRI-eIF2a-ATF4 axis represses globin synthesis under low heme or iron conditions in order to keep excess globins in line with the availability of cofactors (Chen and Zhang, 2019; Crosby et al., 2000; Suragani et al., 2012). Unfolded proteins such as α-chain aggregates can also activate HRI-eIF2a-ATF4, leading to the integrated stress response and repression of protein synthesis. We found that knockout of HBB leads to the formation of α-chain aggregates but paradoxically reduces ATF4 signaling. This suggests that excess heme or iron remaining after loss of *HBB* and concomitant reduction in intact HbA might play a dominant role in reducing ATF4 activity. Reduced ATF4 activity could reactivate globin synthesis, and the formation of HbF would allow coordination of cofactors and a return to homeostasis.

The dramatic increase in γ-globin caused by *ex vivo* reduction in *HBB* raises the question of why β-thalassemia phenotypes are not always ameliorated by ATF4-mediated globin compensation. β-thalassemia patients can express high levels of HbF over long periods of time (Galanello et al., 1989; Manca and Masala, 2008; Rochette et al., 1994), but this phenomenon is not ubiquitous. Mouse models have indicated that the HRI-eIF2a-ATF4 axis is protective against some forms of β-thalassemia, at least in part by upregulating fetal globin (Han et al., 2005; Suragani et al., 2012). This is at odds with observations that knockout of *EIF2AK1* is protective against sickle cell anemia phenotypes by inducing HbF (Grevet et al., 2018), and our own data that knockout of ATF4 or reductions in ATF4 levels downstream of HBB knockout both induce high levels of HbF. Since HRI-eIF2a-ATF4 is involved in the integrated stress response in all cells, reductions in this signaling axis might be negatively selected for during development or otherwise compensated in individuals with β-thalassemia. In this case, developing a cell-type specific approach to disrupting the HRI-eIF2a-ATF4 axis after birth (e.g. during *ex vivo* editing of HSPCs) could be an important avenue for the treatment of hemoglobinopathies. The molecular pathways we find at work during *ex vivo* culture could also be quite different during long term function *in vivo.* We previously observed high levels of HbF in *HBB* edited LT-HSCs recovered after 4 months of xenotransplantation in NBSGW mice (https://www.biorxiv.org/content/10.1101/432716v6), but this does not conclusively address long term function in an individual with β-thalassemia. Determining the long term roles of the HRI-eIF2a-ATF4 axis in patient HSPCs will be an important future step to further unraveling the mechanisms of stress erythropoiesis.

## Methods

### Generation of CRISPRi HUDEP-2 cells

WT-HUDEP2-2 or HBB ko cells were transduced with lentivirus containing the construct EF1a-dCas9-HA-BFP-KRAB-NLS. The cells were FACs sorted for BFP expression and single-cell cloned. Individual clones were transduced with lentivirus containing guide RNAs targeting either *CD55, CD59, or HBB.* The clones were validated for successful knockdown by flow cytometry staining for extracellular markers CD55 and CD59 or intracellularly stained for β-globin.

### Lenti-viral Packaging

Lenti-viral packaging of all constructs was performed using HEK-293T cells. TransIT®-LT1 Transfection Reagent (Mirus) was used following manufacturers instructions. The plasmid mixture contained 50% construct plasmid, 40%DVPR, and 10% VSVG. Viral supernatant was harvested after 48 and 72 hours and filtered through 0.45 μM. For transduction of HUDEP-2 cells, cells were cultured in 50% HUDEP-2 media and 50% viral supernatant for 24 hours.

### sgRNA Plasmid Cloning

sgRNA guide sequences for CRISPRi transcriptional repression were obtained from the Weissman CRISPRi-v2 library (Horlbeck et al., 2016). The chosen guides were cloned into pGL1-library vector (Addgene 84832). All guides used are listed in Supplementary Table 2.

### Cas9 RNP Nucleofection

Cas9 RNP was performed as described previously (Lingeman et al., 2017)

Briefly, IVT guides are purified and complexed with purified Cas9-NLS protein. The nucleofection was performed using Lonza 4D-Nucleofector and using the P3 Primary Cell 96-well NucleofectorTM Kit (V4SP-3096) following manufacturer’s instructions. The HUDEP-2 nucleofector code used was DD-100 and for primary HSPCs ER-100.

### IVT sgRNA

Guide RNAs were in vitro transcribed as described previously (Lingeman et al., 2017). Briefly, guide sequences were ordered as oligonucleotides and formed into duplexes using a PCR thermocycler. The DNA template was transcribed to RNA using HiScribe™ T7 High Yield RNA Synthesis Kit (E2040S) following manufacturer protocol. The resulting RNA was purified using RNeasy Mini kit (74104) and Rnase-Free DnaseI Kit (79254)

### High Pressure Liquid Chromatography and Mass Spectrometry

WT-HUDEP-2 and HBBko cells were differentiated and harvested for lysis in hemolysate reagent containing 0.005M EDTA and 0.07% KCN at 10,000 cells per microliter. The lysis was incubated at room temperature for ten minutes and then centrifuged at max speed for 5 minutes. The supernatant was collected and run on Agilent 1260 Infinity II using a PolyCAT A column, 35×4.6mm (3μm;1500Å) Serial# B19916E; Lot# 16-133-3 3.54CT0315. The following Buffer compositions were used: Mobile Phase A: 20mM Bis-tris, 2mM NaCN pH 6.8 and Mobile Phase B: 20mM Bis-tris, 2mM NaCN, 200mM NaCl, pH 6.9. The following flow settings were used: Gradient: 0-8’ 2-25% Phase B, 8-18’ 25-100% Phase B, 18-23’ 100-2% Mobile Phase B using a Flow Rate: 1.5mL/min and measuring detection of 415nm Diode Array. Three fractions from the HBBko and one fraction from the WT samples were collected and were then processed by the Proteomics group of Functional Genomics Center Zurich (FGCZ) for Proteolytic digestion ZipTip and analysis by LC/MS. Briefly, 100ul of samples were digested with trypsin (5 ng/ul in 10 mM Tris/2 mM CaCl2, pH 8.2) and 2 ul buffer (10 mM Tris/2 mM CaCl2, pH 8.2). Samples were microwaved for 30 minutes at 60C. The samples were dried, dissolved in 20 ul 0.1% TFA and subjected to C18 ZipTip desalting. The eluted sample (10ul of 50% CAN, 0.1%TFA) was dried, dissolved in 20ul of 0.1% FA and transferred to autosampler vials for LC/MS/MS and 1 ul was used for injection.

### HUDEP2 cell culture and differentiation

All cell culture was performed at 37°C in a humidified atmosphere containing 5% CO2. HUDEP-2 cells were cultured in a base medium of SFEM (Stemcell Technologies 9650) containing to a final concentration of dexamethasone 1uM (Sigma D4902-100MG), doxycycline 1ug/ml (Sigma D9891-1G), human stem cell factor 50ng/ml (PeproTech 300-07), erythropoietin 50ng/ml (Peprotech 100-64), and penstrept 1%. Cells were cultured at a density of 2e5 – 1e6 cells/ml. For differentiation, HUDEP-2 cells are centrifuged at 500g for 5 minutes, media is removed and replaced with differentiation media. Differentiation media consists of a base media of IMDM+Glutamax (ThermoFisher 31980030) containing to a final concentration human serum 5% (Sigma H4522-100mL), heparin 2IU/ml (Sigma H3149-25KU), insulin 10ug/ml (Sigma I2643-25mg), erythropoietin 50ng/ml (Peprotech 100-64), holo-transferrin 500ug/ml (Sigma T0665-100mg), mifepristone 1uM (Sigma M8046-100MG), and doxycyline 1ug/ml (Sigma D9891-1G). Cells are differentiated for 5 days and then harvested for analysis.

### mPB-HSPCs cell culture and differentiation

For editing for human CD34+ cells, CD34+ mobilized peripheral blood HSPCs were thawed and cultured in SFEM containing CC110 supplement (Stemcell Technologies 02697) for 2 days. CD34+ cells were then electroporated and recovered for 24 hours in SFEM with CC110. After recovery, cells were transferred into erythroid expansion media containing SFEM and erythroid expansion supplement for 7 days and cultured at a density of 2e5-1e6 cells/ml. The resulting erythroid progenitor cells were transferred to differentiation media containing SFEM with 50ng/ml erythropoietin, 3% normal human serum, and 1 μM mifepristone. Cells were harvested for analysis after 5 days of differentiation.

For editing of human erythroid progenitor cells, CD34+ mobilized peripheral blood HSPCs were thawed and cultured in SFEM containing CC110 supplement (Stemcell Technologies 02697) for 2 days for thaw recovery. Cells are then transferred to erythroid expansion media containing SFEM and erythroid expansion supplement for 5 days and then electroporated. Cells are cultured in erythroid expansion media for another 7 days at a density of 2e5-1e6 cells/ml. The edited erythroid progenitor cells were transferred to differentiation media and harvested for analysis after 5 days of differentiation.

### K562 cell culture

All cell culture was performed at 37°C in a humidified atmosphere containing 5% CO2. K562 cells were grown in a base media of RPMI 1640 GlutaMAX (Gibco 61870010) supplemented with 10% fetal bovine serum, 10% sodium pyruvate, and 1% penstrept. Cells were cultured at a density of 2e5 – 1e6 cells/ml.

### HEK293T cell culture

All cell culture was performed at 37°C in a humidified atmosphere containing 5% CO2. HEK293T cells were grown in a base media of DMEM supplemented with 10% fetal bovine serum, 10% sodium pyruvate, and 1% penstrept. Cells were cultured at a density of 2e5 – 1e6 cells/ml.

### Intracellular FACs Staining

Staining was performed as adapted from the existing methods [Ref Dewitt]. Briefly, undifferentiated or differentiated HUDEP-2 cells were centrifuged at 500g for 5 minutes. The cells were washed with PBS with 0.1% BSA, re-centrifuged, and fixed in 0.05% glutaraldehyde. The fixed cells were centrifuged and washed and re-suspended in PBS with 0.1% BSA and 0.1% Triton-X 100 for permeabilization. The fixed and permeabilized cells were then centrifuged and washed. Antibodies were diluted in PBS with 0.1% BSA and incubated with cells for 20 minutes. The following antibody dilutions were used: Human Fetal Hemoglobin APC (Thermo Fisher MHFH05) 1:10, Anti-Human Fetal Hemoglobin FITC 1:10 (BD Pharmigen 552829), and Anti-Hemoglobin B-(37-8) PE/FITC 1:100 (Santa Cruz Biotechnology SC-21757 PE, SC-21757 FITC). After staining, the cells were centrifuged and washed twice before analysis by flow cytometry.

### RNA-SEQ

WT-HUDEP-2 and HBBko cells were cultured and differentiated in triplicates. Cells were harvested and RNA was extracted using the RNeasy Mini kit 74104, Rnase-Free DnaseI Kit 79254 and Qiashredder 79654. RNA concentrations were quantified using Qubit™ RNA BR Assay (ThermoFisher) and 500ng were used for library preparation. RNA-seq library was made using Illumina TruSeq RNA Library Prep Kit v2 and following manufacturer’s instructions. Reads for all eighteen samples (three replicates of HBBko and three WT HUDEP-2 samples at three time-points) were quantified using kallisto [PMID: 27043002] and the hg38 index to assign reads to transcripts. Differential analysis of transcript abundance and consolidation of individual transcripts to gene-level abundance was calculated using sleuth [PMID: 28581496].

### qRT-PCR

RNA was harvested from cells using Qiagen RNeasy Mini Kit and Rnase-Free DnaseI Kit following manufacturer’s instructions. RNA was reverse transcribed to cDNA using Iscript™ Reverse Transcription Supermix (BioRad) and qRT-PCR reactions were set up using SsoAdvanced Universal SYBR Green or SsoFast™ EvaGreen Supermix (BioRad). Reactions were run on the StepOne Plus Real-Time PCR System (Applied Biosystems) or the QuantStudio 6 Flex (Thermo Fisher). Samples were analyzed using a two-step amplification and melt curves were obtained after 40 cycles. The Ct values for genes of interest were normalized to GAPDH, and expressions of genes are represented as 2-[ΔCt] or 2-[ΔΔCt] for fold change over control condition. All primers used for qRT-PCR are listed in Table S2.

### ChIP-qPCR

ChIP was performed as described in (Wienert et al., 2019). Briefly, 10 million cells per sample were harvested and cross-linked in 1% Formaldehyde. Cross-linking was quenched with the addition of 1.5M glycine. Samples were then lysed for 10 minutes at 4C in 50 mM Hepes–KOH, pH 7.5; 140 mM NaCl; 1 mM EDTA; 10% glycerol; 0.5% NP-40 or Igepal CA-630; 0.25% Triton X-100. Cells were then centrifuged at 1500g for 3 minutes and the supernatant was discarded. The pellet was resuspended in 10 mM Tris–HCl, pH8.0; 200 mM NaCl; 1 mM EDTA; 0.5 mM EGTA and incubated for 5 minutes at 4C. The cells were then centrifuged at 1500g for 3 minutes and the supernatant was discarded. THe pellet was resuspended in 10 mM Tris–HCl, pH 8; 100 mM NaCl; 1 mM EDTA; 0.5 mM EGTA; 0.1% Na–Deoxycholate; 0.5% N-lauroylsarcosine and sonicated using the Covaris S220 following manufacturer’s instructions. Protein A beads (ThermoFisher) were complexed with antibody and the antibody-bead complexes were incubated with cell lysates at 4C overnight with rotation. The antibodies used were rabbit anti-ATF4 (CST 11815S) and rabbit IgG (Novus Biologicals NBP2-24891). The beads were retrieved using a magnetic stand and rinsed with RIPA buffer. Elution buffer containing 50 mM Tris–HCl, pH 8; 10 mM EDTA; 1% SDS was added to the beads for reverse crosslinking at 65C overnight with shaking. After reverse crosslinking, the beads were removed. The eluted DNA was treated with RNaseA and Proteinase K and then purified using Qiagen MinElute PCR Purification Kit, following the manufacturer’s instructions. Q-PCR reactions were set up using SsoAdvanced Universal SYBR Green or SsoFast™ EvaGreen Supermix (BioRad). Reactions were run on the StepOne Plus Real-Time PCR System (Applied Biosystems) or the QuantStudio 6 Flex (Thermo Fisher). The Ct values were analyzed by the enrichment compared to input method.

### ChIP-seq

ChIP was performed as described in ChIP-qPCR method section. Sequencing library was prepared using NEBNext Ultra II DNA Library Prep Kit for Illumina (E7647) and NEBNext Multiplex Oligos for Illumina (Dual Index Primers Set 1) following manufacturer’s instructions. Paired-end 150bp reads were generated on an Illumina NextSeq500 at the Functional Genomics Center Zürich (FGCZ) and demultiplexed. FastQC (Andrews S. (2010) A quality control tool for high throughput sequence data. http://www.bioinformatics.babraham.ac.uk/projects/fastqc/) was used for initial quality control of reads. All samples, WT diff and undiff, HBBko diff and undiff, ATF4KO3 and IgG, were processed according to ENCODE guidelines for unreplicated transcription factor ChIP-seq analysis (Landt et al., 2012). In detail, raw reads were aligned against GRCh38 using bowtie2. Duplicate reads were marked using Picard’s MarkDuplicates and multimapping, low quality, duplicated and non-properly paired reads were removed. Library complexity measures and flagstats were generated for each BAM file. BAM files were converted to tagAlign format and two subsampled pseudoreplicates were generated for each sample with half the total reads. Peak calling, fold change and p-value signal tracks were generated using MACS2 (Zhang et al., 2008). Irreproducible Discovery Rate (IDR) analysis was performed using self-pseudoreplicates and the main samples to obtain self-consistent sets of peaks. Final peak calls were filtered by ENCODE blacklist (Amemiya et al., 2019) and an IDR of 2%.

Sets of peaks for each comparison were analysed and associated to genes using the R package ChIPseeker (Yu et al., 2015) and Bioconductor hg38 TxDb (Team BC, Maintainer BP (2019). TxDb.Hsapiens.UCSC.hg38.knownGene: Annotation package for TxDb object(s). R package version 3.4.6.). ChIP-seq peaks and RNA-seq results were merged using HGNC symbols using custom scripts. Figures were generated using the Integrative Genome Viewer (Robinson et al., 2011) and the UCSC Genome Brower (Kent et al., 2002).

### Western Blot

Cells were harvested and lysed using RIPA buffer (Millipore) supplemented with Halt Protease Inhibitor Cocktail (ThermoFisher) and Phosphatase Inhibitor (ThermoFisher). Cell lysate concentrations were measured using Bradford reagent (VWR) or BCA assay (ThermoFisher). Cell lysates were normalized using NuPage LDS 4x Sample Buffer (Invitrogen) and samples were run on NU-PAGE Noves Bis-Tris 4–12% gels (Invitrogen) in NuPAGE™ MES SDS Running Buffer at 180V for 40 minutes. Protein gels were transferred to a 0.4-μm nitrocellulose membranes at 1.3 A and 25 V for 15 to 20 minutes in a semi-dry apparatus (BioRad). After protein transfer, membranes were blocked in Tris-buffered saline with 1% Tween-20 (TBS-T) containing 5% nonfat milk for 30 minutes at room temperature with rocking. Primary antibodies were diluted in TBST containing 5%BSA and 0.1% sodium azide. The following antibody concentrations were used: rabbit anti-ATF4 (CST 11815S) 1:1000, mouse anti-BCL11A (Abcam ab19489) 1:1000, rabbit anti-LRF (ThermoFisher PA528144) 1:1000, mouse anti-GAPDH D4C6R (CST 2118S) 1:1000, Rabbit anti-GAPDH 14C10 (CST 2118S) 1:1000, Mouse anti-B-Globin (SCBT sc-21757) 1:500, Rabbit anti-Fetal-Globin (Abcam ab137096) 1:1000.

Membranes were rinsed in TBST and incubated with antibody for 2 hours at room temperature or 4C overnight with rocking. After incubation, membranes are rinsed twice with TBST for 5 minutes. Secondary antibodies are diluted in 1% Tween-20 (TBS-T) containing 5% nonfat milk and incubated at room temperature for 30 minutes with rocking. The secondary antibodies used for Western blotting were obtained from Li-Cor and are as follows: donkey anti-mouse IRDye 680CW (926-32222), donkey anti-mouse IRDye 800CW (926-32212), donkey anti-rabbit IRDye 680CW (926-32223), and donkey anti-rabbit IRDye 800CW (926-32213). Membranes are then rinsed twice with TBST for 5 minutes each. Membranes are rinsed with PBS for 5 minutes and then imaged using Li-Cor Odyssey CLx.

## References

Alter, B.P. (1979). Fetal erythropoiesis in stress hematopoiesis. Exp. Hematol. 7 Suppl 5, 200–209.

Amemiya, H.M., Kundaje, A., and Boyle, A.P. (2019). The ENCODE Blacklist: Identification of Problematic Regions of the Genome. Sci. Rep. 9, 1–5.

Bauer, D.E., Kamran, S.C., Lessard, S., Xu, J., Fujiwara, Y., Lin, C., Shao, Z., Canver, M.C., Smith, E.C., Pinello, L., et al. (2013). An erythroid enhancer of BCL11A subject to genetic variation determines fetal hemoglobin level. Sci. N. Y. NY 342, 253–257.

Berry, M., Grosveld, F., and Dillon, N. (1992). A single point mutation is the cause of the Greek form of hereditary persistence of fetal haemoglobin. Nature 358, 499–502.

Chen, J.-J., and Zhang, S. (2019). Heme-regulated eIF2α kinase in erythropoiesis and hemoglobinopathies. Blood 134, 1697–1707.

Chung, J.E., Magis, W., Vu, J., Heo, S.-J., Wartiovaara, K., Walters, M.C., Kurita, R., Nakamura, Y., Boffelli, D., Martin, D.I.K., et al. (2019). CRISPR-Cas9 interrogation of a putative fetal globin repressor in human erythroid cells. PLOS ONE 14, e0208237.

Crosby, J.S., Chefalo, P.J., Yeh, I., Ying, S., London, I.M., Leboulch, P., and Chen, J.J. (2000). Regulation of hemoglobin synthesis and proliferation of differentiating erythroid cells by heme-regulated eIF-2alpha kinase. Blood 96, 3241–3248.

DeWitt, M.A., Magis, W., Bray, N.L., Wang, T., Berman, J.R., Urbinati, F., Heo, S.-J., Mitros, T., Muñoz, D.P., Boffelli, D., et al. (2016). Selection-free genome editing of the sickle mutation in human adult hematopoietic stem/progenitor cells. Sci. Transl. Med. 8, 360ra134–360ra134.

El-Brolosy, M.A., Kontarakis, Z., Rossi, A., Kuenne, C., Günther, S., Fukuda, N., Kikhi, K., Boezio, G.L.M., Takacs, C.M., Lai, S.-L., et al. (2019). Genetic compensation triggered by mutant mRNA degradation. Nature 568, 193–197.

Galanello, R., Barella, S., Maccioni, L., Paglietti, E., Melis, M.A., Rosatelli, M.C., Argiolu, F., and Cao, A. (1989). Erythropoiesis following bone marrow transplantation from donors heterozygous for ß-thalassaemia. Br. J. Haematol. 72, 561–566.

Grevet, J.D., Lan, X., Hamagami, N., Edwards, C.R., Sankaranarayanan, L., Ji, X., Bhardwaj, S.K., Face, C.J., Posocco, D.F., Abdulmalik, O., et al. (2018). Domain-focused CRISPR screen identifies HRI as a fetal hemoglobin regulator in human erythroid cells. Science 361, 285–290.

Han, A.-P., Fleming, M.D., and Chen, J.-J. (2005). Heme-regulated eIF2α kinase modifies the phenotypic severity of murine models of erythropoietic protoporphyria and ß-thalassemia. J. Clin. Invest. 115, 1562–1570.

Huang, D.W., Sherman, B.T., and Lempicki, R.A. (2009). Systematic and integrative analysis of large gene lists using DAVID bioinformatics resources. Nat. Protoc. 4, 44–57.

Jacob, G.F., and Raper, A.B. (1958). Hereditary Persistence of Foetal Haemoglobin Production, and its Interaction with the Sickle-Cell Trait. Br. J. Haematol. 4, 138–149.

Kent, W.J., Sugnet, C.W., Furey, T.S., Roskin, K.M., Pringle, T.H., Zahler, A.M., and Haussler, and D. (2002). The Human Genome Browser at UCSC. Genome Res. 12, 996–1006.

Krämer, A., Green, J., Pollard, J., and Tugendreich, S. (2014). Causal analysis approaches in Ingenuity Pathway Analysis. Bioinformatics 30, 523–530.

Kuleshov, M.V., Jones, M.R., Rouillard, A.D., Fernandez, N.F., Duan, Q., Wang, Z., Koplev, S., Jenkins, S.L., Jagodnik, K.M., Lachmann, A., et al. (2016). Enrichr: a comprehensive gene set enrichment analysis web server 2016 update. Nucleic Acids Res. 44, W90–W97.

Kurita, R., Suda, N., Sudo, K., Miharada, K., Hiroyama, T., Miyoshi, H., Tani, K., and Nakamura, Y. (2013). Establishment of immortalized human erythroid progenitor cell lines able to produce enucleated red blood cells. PloS One 8, e59890.

Landt, S.G., Marinov, G.K., Kundaje, A., Kheradpour, P., Pauli, F., Batzoglou, S., Bernstein, B. E., Bickel, P., Brown, J.B., Cayting, P., et al. (2012). ChIP-seq guidelines and practices of the ENCODE and modENCODE consortia. Genome Res. 22, 1813–1831.

Lechauve, C., Keith, J., Khandros, E., Fowler, S., Mayberry, K., Freiwan, A., Thom, C.S., Delbini, P., Romero, E.B., Zhang, J., et al. (2019). The autophagy-activating kinase ULK1 mediates clearance of free a-globin in ß-thalassemia. Sci. Transl. Med. 11, eaav4881.

Lingeman, E., Jeans, C., and Corn, J.E. (2017). Production of Purified CasRNPs for Efficacious Genome Editing. Curr. Protoc. Mol. Biol. Ed. Frederick M Ausubel Al 120, 31.10.1–31.10.19.

Ma, Z., Zhu, P., Shi, H., Guo, L., Zhang, Q., Chen, Y., Chen, S., Zhang, Z., Peng, J., and Chen, J. (2019). PTC-bearing mRNA elicits a genetic compensation response via Upf3a and COMPASS components. Nature 568, 259–263.

Manca, L., and Masala, B. (2008). Disorders of the synthesis of human fetal hemoglobin. IUBMB Life 60, 94–111.

Masuoka, H.C., and Townes, T.M. (2002). Targeted disruption of the activating transcription factor 4 gene results in severe fetal anemia in mice. Blood 99, 736–745.

Meletis, J., Papavasiliou, S., Yataganas, X., Vavourakis, S., Konstantopoulos, K., Poziopoulos, C., Samarkos, M., Michali, E., Dalekou, M., and Eliopoulos, G. (1994). “Fetal” erythropoiesis following bone marrow transplantation as estimated by the number of F cells in the peripheral blood. Bone Marrow Transplant. 14, 737–740.

Papayannopoulou, T., Vichinsky, E., and Stamatoyannopoulos, G. (1980). Fetal Hb Production during Acute Erythroid Expansion: I. OBSERVATIONS IN PATIENTS WITH TRANSIENT ERYTHROBLASTOPENIA AND POST-PHLEBOTOMY. Br. J. Haematol. 44, 535–546.

Platt, O.S., Orkin, S.H., Dover, G., Beardsley, G.P., Miller, B., and Nathan, D.G. (1984). Hydroxyurea enhances fetal hemoglobin production in sickle cell anemia. J. Clin. Invest. 74, 652–656.

Robinson, J.T., Thorvaldsdóttir, H., Winckler, W., Guttman, M., Lander, E.S., Getz, G., and Mesirov, J.P. (2011). Integrative Genomics Viewer. Nat. Biotechnol. 29, 24–26.

Rochette, J., Craig, J.E., Thein, S.L., and Rochette, J. (1994). Fetal hemoglobin levels in adults. Blood Rev. 8, 213–224.

Rossi, A., Kontarakis, Z., Gerri, C., Nolte, H., Hölper, S., Krüger, M., and Stainier, D.Y.R. (2015). Genetic compensation induced by deleterious mutations but not gene knockdowns. Nature 524, 230–233.

Sankaran, V.G. (2011). Targeted Therapeutic Strategies for Fetal Hemoglobin Induction. Hematology 2011, 459–465.

Schiroli, G., Conti, A., Ferrari, S., Della Volpe, L., Jacob, A., Albano, L., Beretta, S., Calabria, A., Vavassori, V., Gasparini, P., et al. (2019). Precise Gene Editing Preserves Hematopoietic Stem Cell Function following Transient p53-Mediated DNA Damage Response. Cell Stem Cell 24, 551–565.e8.

Steinmüller, L., and Thiel, G. (2005). Regulation of Gene Transcription by a Constitutively Active Mutant of Activating Transcription Factor 2 (ATF2). Biol. Chem. 384, 667–672.

Suragani, R.N.V.S., Zachariah, R.S., Velazquez, J.G., Liu, S., Sun, C.-W., Townes, T.M., and Chen, J.-J. (2012). Heme-regulated eIF2α kinase activated Atf4 signaling pathway in oxidative stress and erythropoiesis. Blood 119, 5276–5284.

Weinberg, R.S., Schofield, J.M., Lenes, A.L., Brochstein, J., and Alter, B.P. (1986). Adult’fetal-like’erythropoiesis characterizes recovery from bone marrow transplantation. Br. J. Haematol. 63, 415–424.

Wienert, B., Funnell, A.P.W., Norton, L.J., Pearson, R.C.M., Wilkinson-White, L.E., Lester, K., Vadolas, J., Porteus, M.H., Matthews, J.M., Quinlan, K.G.R., et al. (2015). Editing the genome to introduce a beneficial naturally occurring mutation associated with increased fetal globin. Nat. Commun. 6, 70–85.

Wienert, B., Martyn, G.E., Funnell, A.P.W., Quinlan, K.G.R., and Crossley, M. (2018). Wake-up Sleepy Gene: Reactivating Fetal Globin for ß-Hemoglobinopathies. Trends Genet. 34, 927–940.

Wienert, B., Wyman, S.K., Richardson, C.D., Yeh, C.D., Akcakaya, P., Porritt, M.J., Morlock, M., Vu, J.T., Kazane, K.R., Watry, H.L., et al. (2019). Unbiased detection of CRISPR off-targets in vivo using DISCOVER-Seq. Science 364, 286–289.

Xiang, J., Wu, D.-C., Chen, Y., and Paulson, R.F. (2015). In vitro culture of stress erythroid progenitors identifies distinct progenitor populations and analogous human progenitors. Blood 125, 1803–1812.

Yu, G., Wang, L.-G., and He, Q.-Y. (2015). ChIPseeker: an R/Bioconductor package for ChIP peak annotation, comparison and visualization. Bioinformatics 31, 2382–2383.

Zhang, S., Macias-Garcia, A., Ulirsch, J.C., Velazquez, J., Butty, V.L., Levine, S.S., Sankaran, V.G., and Chen, J.-J. (2019). HRI coordinates translation necessary for protein homeostasis and mitochondrial function in erythropoiesis. ELife 8, e46976.

Zhang, Y., Liu, T., Meyer, C.A., Eeckhoute, J., Johnson, D.S., Bernstein, B.E., Nusbaum, C., Myers, R.M., Brown, M., Li, W., et al. (2008). Model-based analysis of ChIP-Seq (MACS). Genome Biol. 9, R137.

