## Supplemental Figure Legend for "ATF4 mediates fetal globin upregulation in response to reduced β-globin"

**Supplement Figure 1 (related to Figure 1)**

A) Schematic showing the e66 (g5) guide location in *HBB*, with the cut site indicated by a black line. The HBBko clone was genotyped by TA cloning of the *HBB* locus followed by Sanger sequencing. Two alleles were found: a 5- and 7-basepair deletion.

B) Flow cytometry plots showing forward and side scattering of WT and HBBko over the course of 5 days in differentiation media. Cells were stained for CD235a (Glycophorin A) and shown as dot plots and overlaid histograms to compare WT to HBBko during differentiation in red and blue, respectively. IgG isotype control trace is in gray.

C) qRT-PCR of *HBB* in undifferentiated and differentiated cells comparing transcript expression levels between WT and HBBko cells.

D) Editing efficiency of the panel of HBB guides in HUDEP-2 cells as measured by TIDE analysis.

E) Mobilized peripheral blood CD34+ cells from a healthy donor were edited with a subset of the HBB guides, recovered in erythroid expansion media, differentiated for 5 days, and analyzed for qRT-PCR for *HBB* and *HBG 1/2* for (n=2) technical replicates for one healthy donor. Editing efficiency was measured by TIDE

F) Editing of human primary erythroid progenitors from another healthy donor is shown in Data is shown for one replicate.

G) Total cell counts of edited CD34+ cells during recovery in erythroid expansion media. Data is shown as the mean ± SEM of (n=2) technical replicates for one healthy donor.

H) Total cell counts of edited CD34+ cells after 5 days of differentiation. Data is shown as the mean ± SD of (n=2) technical replicates for one healthy donor.

I) HBBko CRISPRi cell line was validated using guides targeting CD55 and CD59. FACs staining was used for validating knockdown of the cell marker. WT-CRISPRi cell line was validated using an HBB guide and differentiated and intracellularly stained for B-globin protein and compared to non-targeting guide.

J) qRT-PCR of *HBB* and *HBG1/2* for CRISPRi knockdown of *HBB* in WT differentiated cells. Data is shown for one replicate.

K) qRT-PCR of HBB and HBG1/2 from pooled knockout of *HBG1/2* in HUDEP-1 cells. N=2 technical replicates are shown as mean ± SD.

**Supplement Figure 2 (related to Figure 2)**

A) MA plots of RNAseq data for undifferentiated cells and 2 day differentiation, with globin genes highlighted.

B) Pearson correlation matrix of RNAseq samples, highlighting inter-condition reproducibility and differences between conditions.

C) Pearson correlation analysis of RNAseq data comparing WT to HBBko at three different time points of differentiation

D) Leftmost panel shows quantified intracellular flow staining of γ-globin for HBBko CRISPRi cells expressing either non-targeting, HBG1/2, SMG6, or XRN1 guide. Right three panels show qRT-PCR for SMG6, XRN1, or HBG1/2 for knockdown validation.

**Supplement Figure 3 (related to Figure 3)**

A) Schematic of ATF4 with sgRNAs used for generation of ATF4 knockouts and ATF4ΔN. See **Table S2** for all guide sequences.

B) PCR genotyping of ATF4ko-1 and ATF4ko-2 clones. Top panel shows PCR amplification using primers outside the cut region. Successful *ATF4* deletion is indicated by ~500 base pair fragment. Unsuccessful deletion results in failure of PCR amplification due to the large size. Lower panel shows PCR amplification of a region within the *ATF4* gene. Successful deletion of *ATF4* is indicated by no amplification.

C) Western blotting for ATF4 of WT and ATF4 mutant clones treated with 40nM cyclopiazonic acid (CPA) for 16 hours. ATF4ΔN removes the amino terminal regulatory region but retains the C-terminal DNA binding domain.

D) Flow cytometry plots of forward and side scatter 5 days post differentiation of WT, HBBko, ATF4ΔN, ATF4ko-1, and ATF4ko-2. Right panel shows percentages of differentiated cells as the mean ± SD of 3 biological replicates.

E) PCR genotyping of ATF4ΔN using primers flanking the ATF4ΔN cut sites. TA-cloning and Sanger sequencing revealed one 398- and one 415-basepair deletion for the ATF4ΔN clone.

F) ATF4 CHIP-qPCR for the *ASNS* promoter region shows ATF4 binding at WT and ATF4ΔN and loss of binding for ATF4ko-1 and ATF4ko-2 in undifferentiated cells. The data is shown as one replicate. Right panel shows qRT-PCR on differentiated cells for *ASNS*. The data is shown as the mean ± SD of 3 biological replicates.

G) Quantification of intracellular FACs staining of differentiated WT, HBBko, and ATF4ΔN cells. Data is presented as mean ± SD of 3 biological replicates.

H) Quantification of intracellular FACs staining of undifferentiated WT, HBBko, ATF4ΔN, and ATF4ko cells. Data is presented as mean ± SD of 4 biological replicates.

I) qRT-PCR of globin gene expression for differentiated WT, HBBko, and ATF4ΔN. Data is presented as mean ± SD of 3 biological replicates.

J) qRT-PCR of *ATF4* and *HBG1/2* in K562 WT cells and two K562 ATF4 knockout clones.

**Supplement Figure 4 (related to Figure 4)**

A) *ATF4, BCL11A*, and *ZBTB7A* transcript levels in HBBko HUDEP-2 cells are shown as log2 fold change relative to WT cells undifferentiated, after 2 days of differentiation, and after 5 days of differentiation.

B) Top panel shows intracellular FACs staining of differentiated cells for HbA and HbF in WT CRISPRi cells with guides against non-targeting, BCL11A, and HBB. Lower panel shows quantified data as mean ± SD of 3 biological replicates.

C) Western blot of undifferentiated WT, HBBko, ATF4ΔN, and ATF4ko cells.

D) Western blot of K562 WT, ATF4ko.562-1 and , ATF4ko.562-2 for BCL11A expression.

E) ATF4 ChIP-seq signal enrichment in WT HUDEP-2 cells and corresponding RNA-seq log2 TPM values.

**Supplement Figure 5 (related to Figure 4)**

A) ATF4 ChIP-seq enrichment tracks for the globin locus. DNAse signal and H3k27ac tracks show reference ENCODE data from K562 cells.
B) ATF4 ChIP-seq fold enrichment tracks for the BCL11A locus.
C) ATF4 ChIP-seq fold enrichment tracks for the ZBTB7A locus.

**Supplement Figure 6**

A) CHAC1 raw read coverage for ATF4 ChIP-seq. Tracks show library size normalized tags per million (TPM) coverage of WT undiff and diff, HBBko undiff and diff, ATF4KO3 undiff and IgG data. DNase signal and H3k27ac tracks show reference ENCODE data from K562 cells. Annotation tracks show GENCODE v32 gene annotation and transcription start site annotation from refTSS v3.1
B) C16orf72 raw read coverage for ATF4 ChIP-seq.
C) IGF2R1 raw read coverage for ATF4 ChIP-seq.

**Supplement Figure 7**

A) Transcript levels of UPR targets in HBBko HUDEP-2 cells are shown as log2 fold change relative to WT cells undifferentiated, after 2 days of differentiation, and after 5 days of differentiation.

**Supplemental Table 1: Mass Spectrometry of HPLC fractions**

Exclusive Spectrum Counts identifying proteins within the 4 fractions collected from HPLC runs. The first 3 fractions correspond to the 3 primary peaks in the differentiated HBBko cells. The fourth fraction was the primary peak from the different WT cells.

**Supplemental Table 2: sgRNAs and oligos**

All sequences used for sgRNA IVT and sgRNAs cloned for for CRISPRi repression. All oligos used for qRT-PCR, ChIP-qpcr, TIDE analysis, and validation of ATF4 knockouts are also listed.
