## Supplemental Figures for "ATF4 mediates fetal globin upregulation in response to reduced β-globin"

### Supplement Fig 1

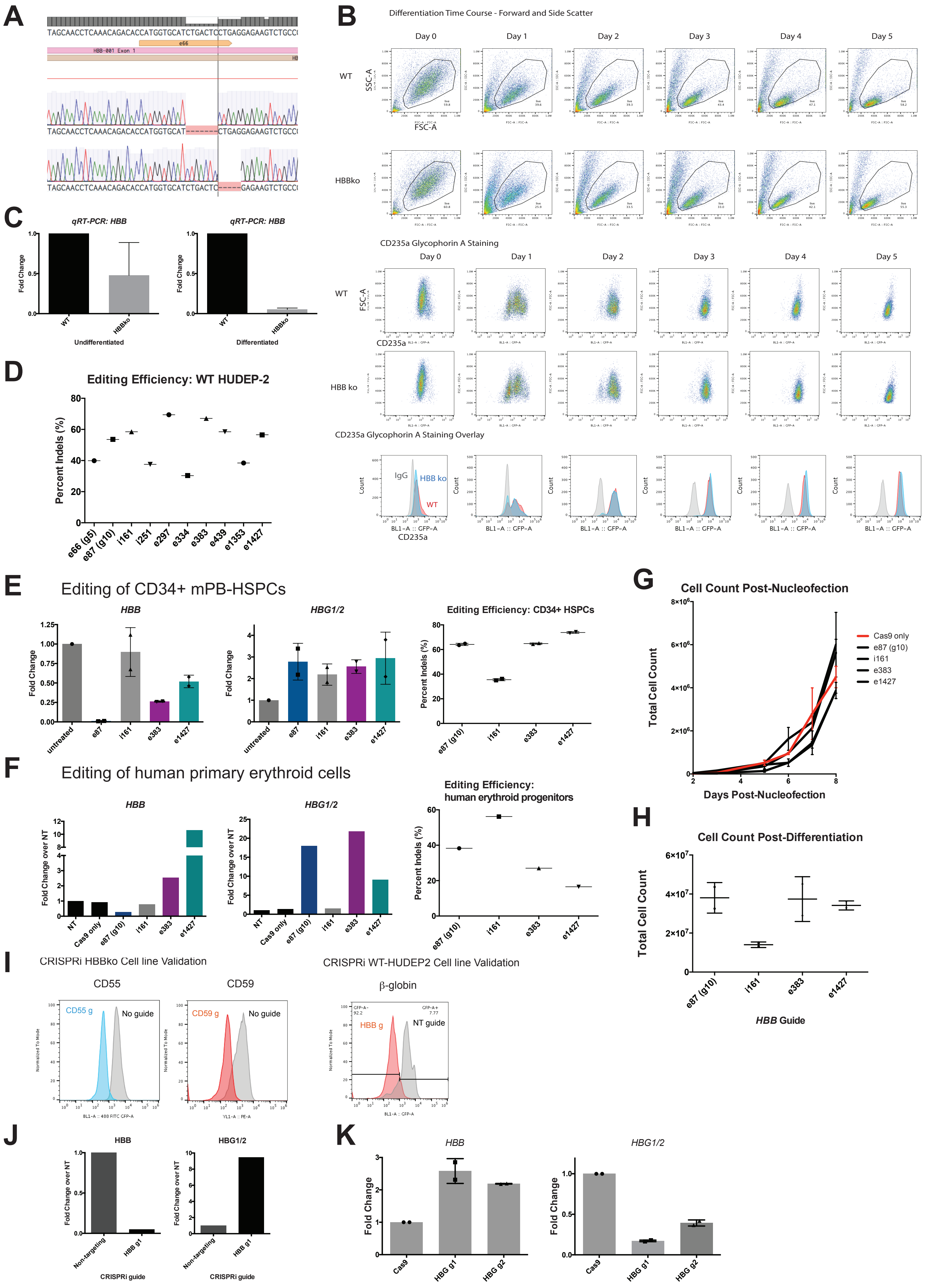

Supplemental Figure 2

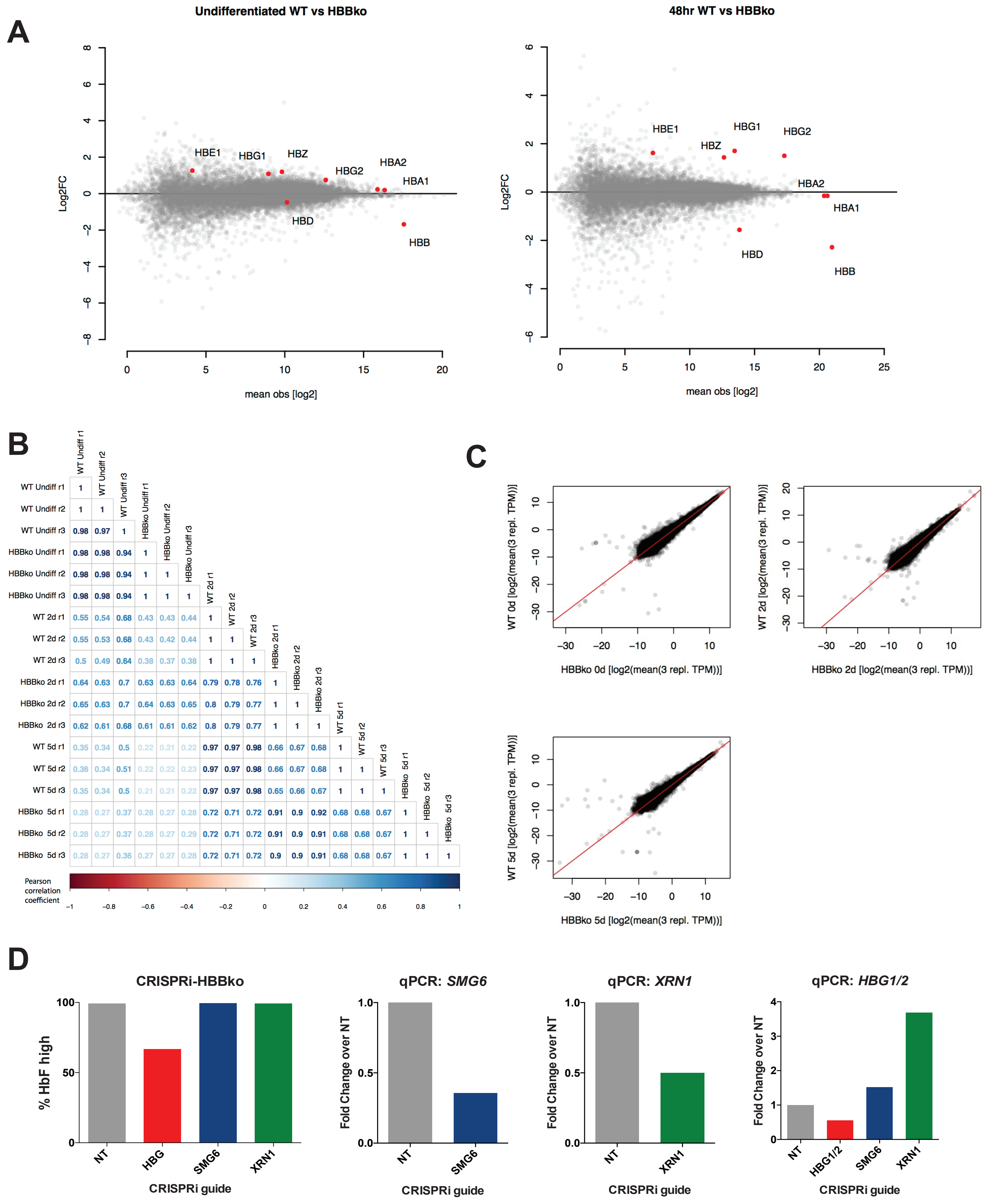

Supplement Figure 3

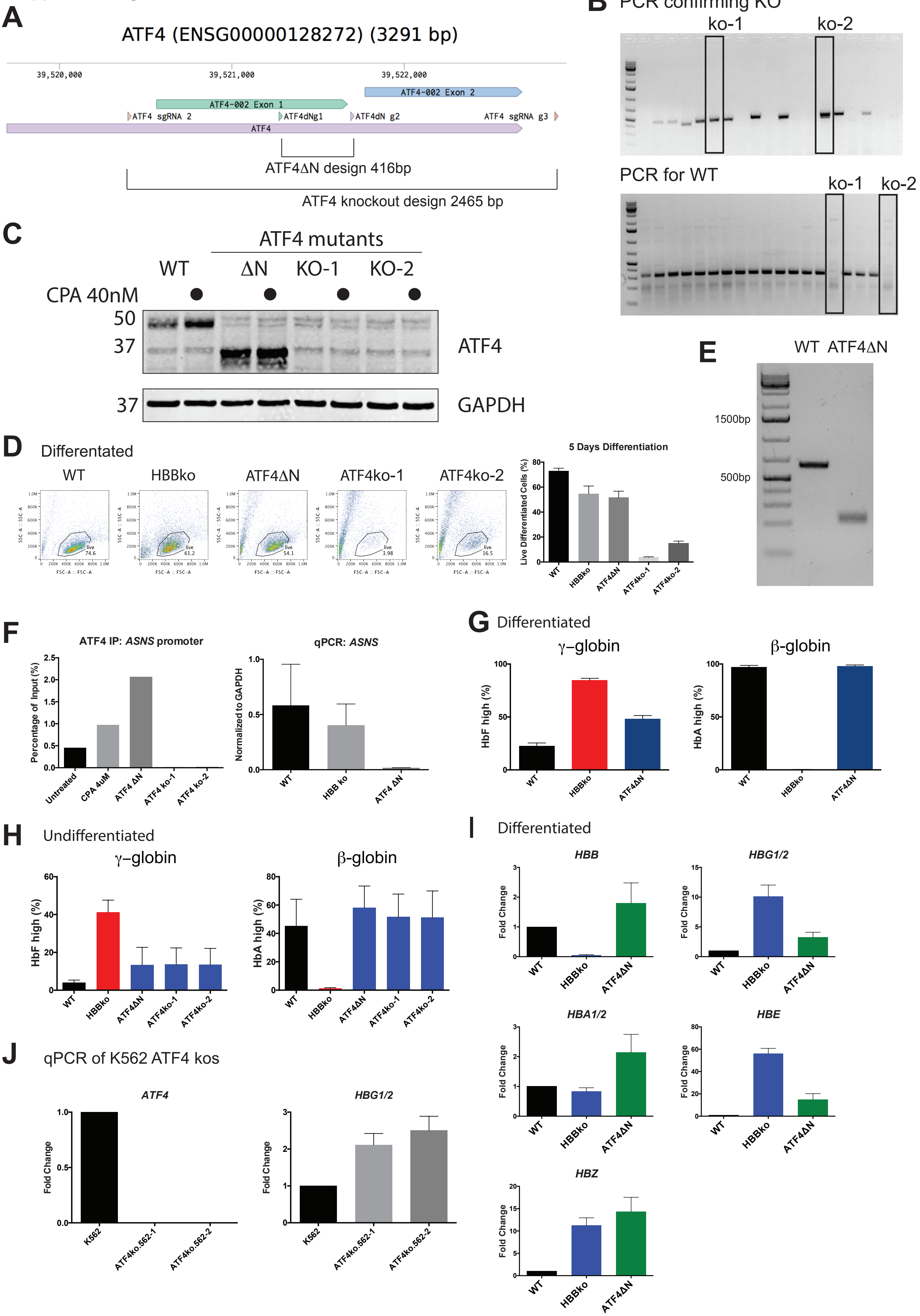

Supplement Fig 4

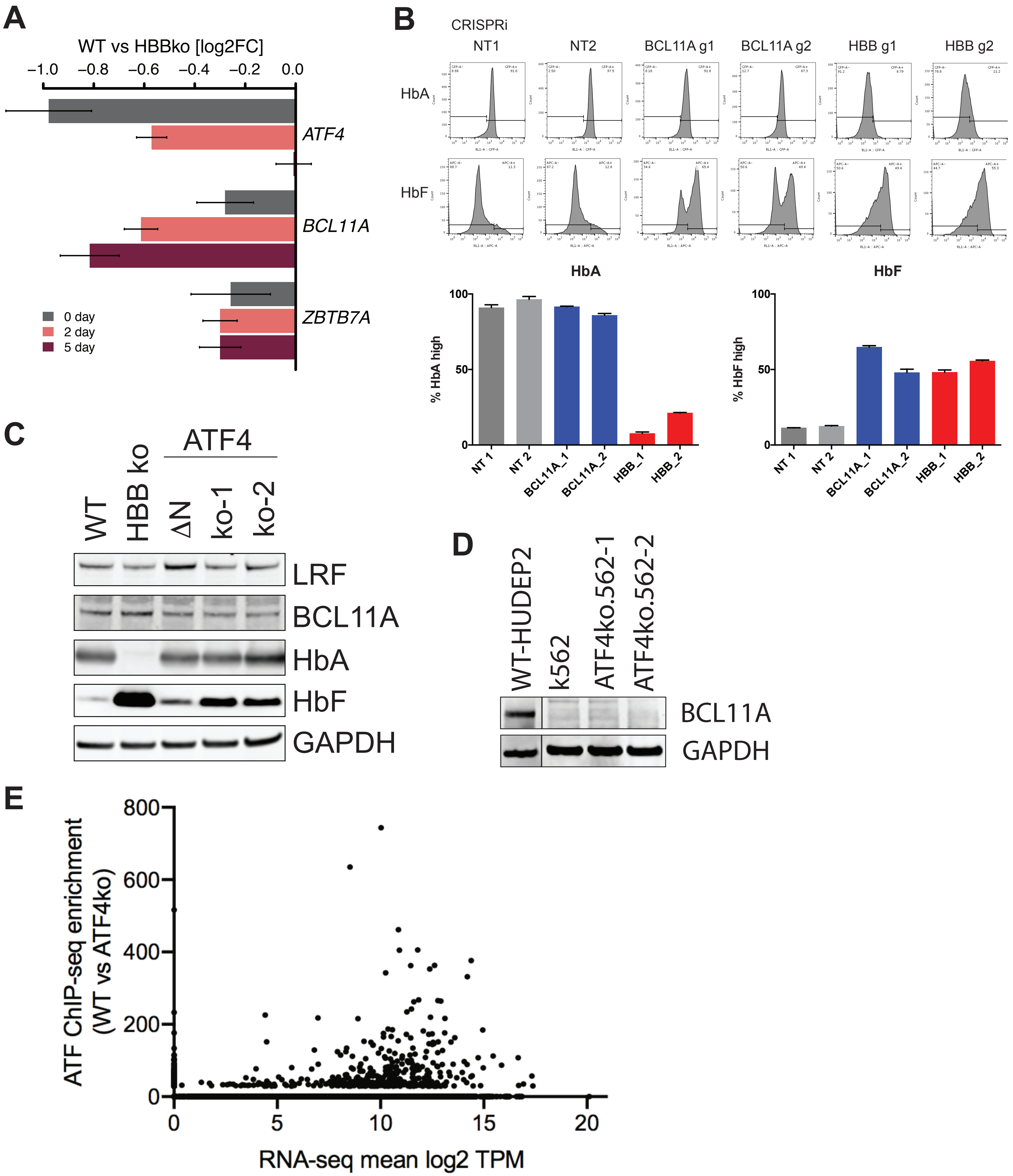

Supplement Fig 5

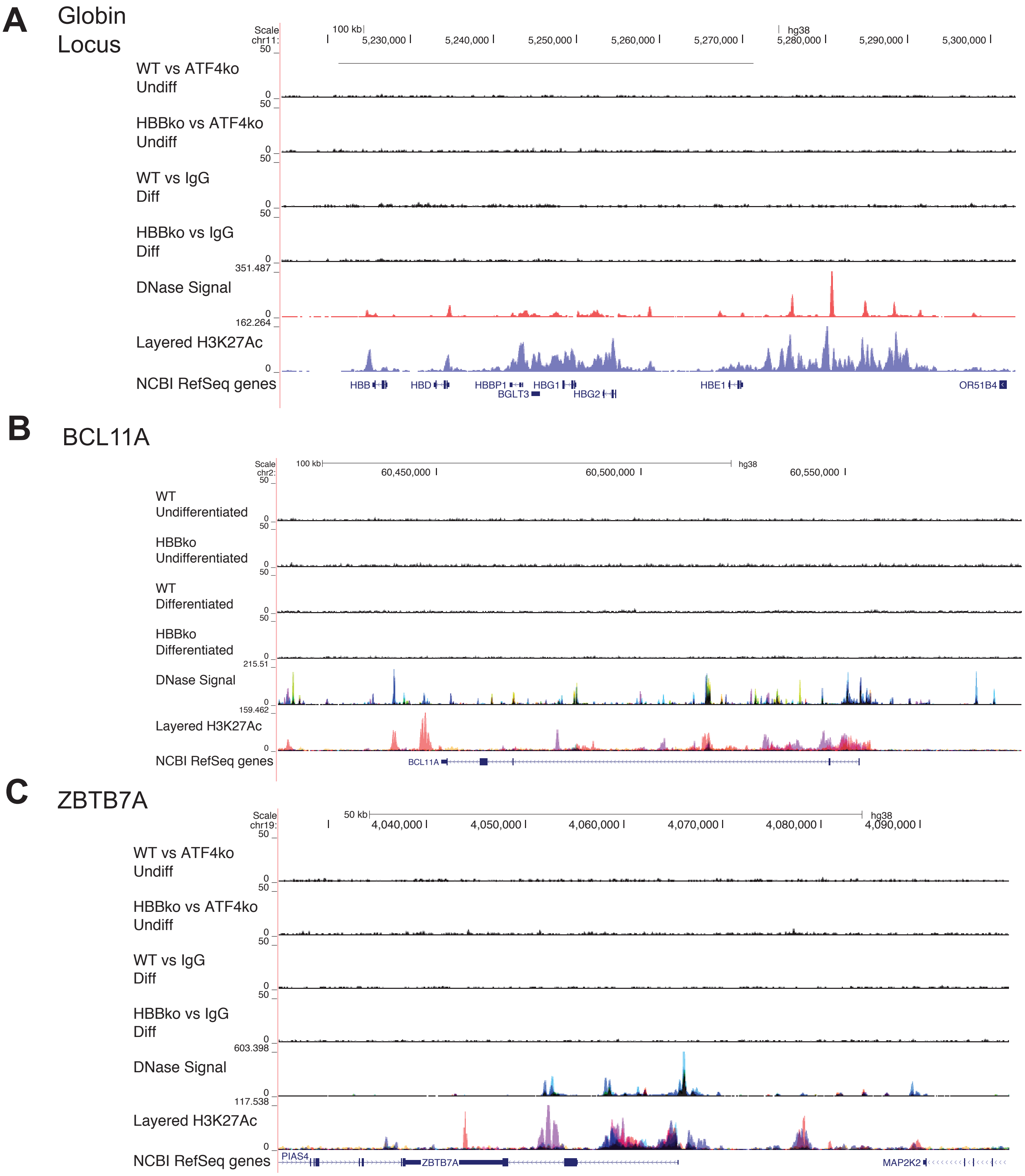

Supplement Fig 6

**A** CHAC1

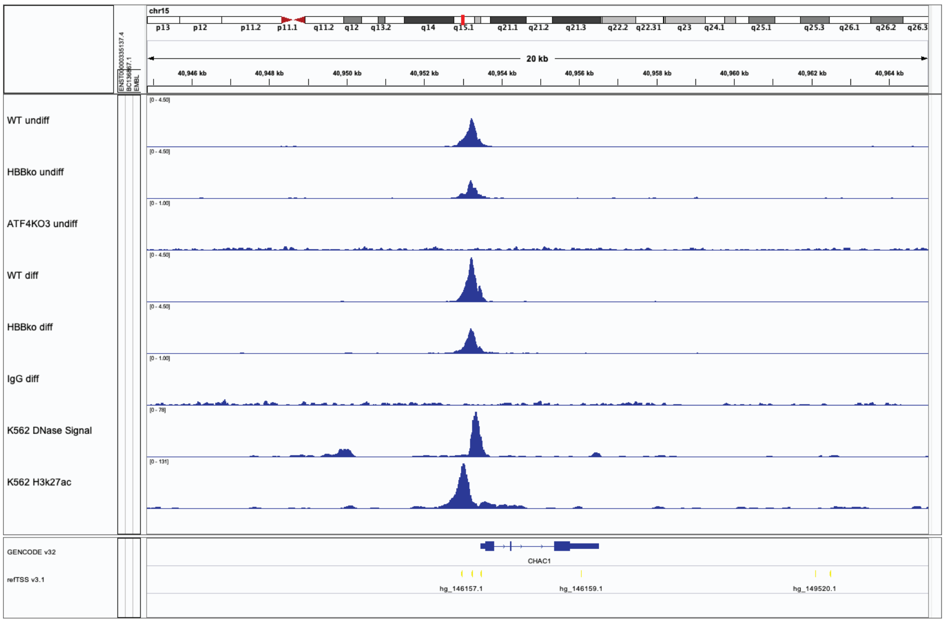


**B** C16orf72

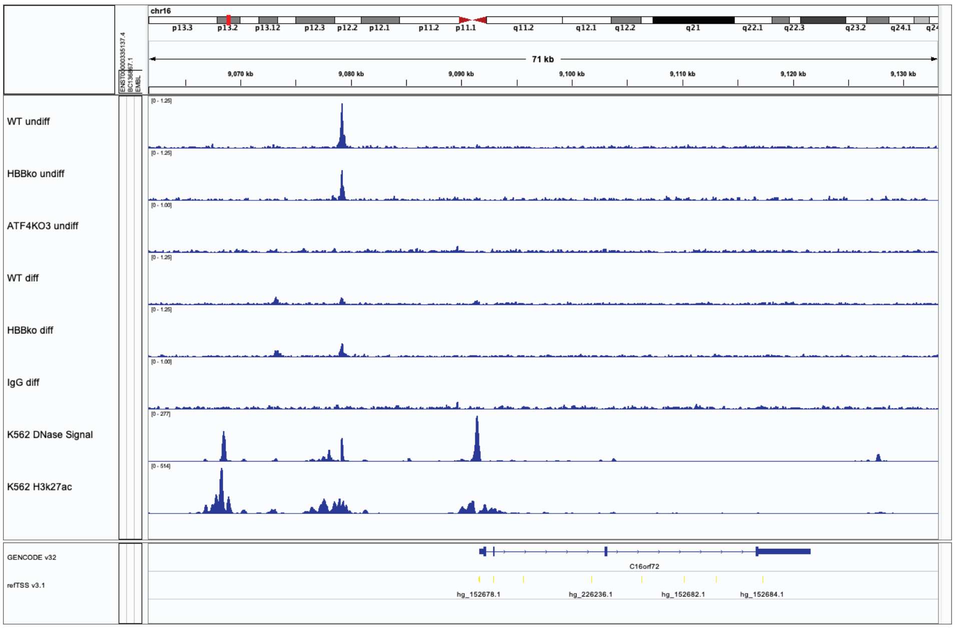


**C** IGF2R

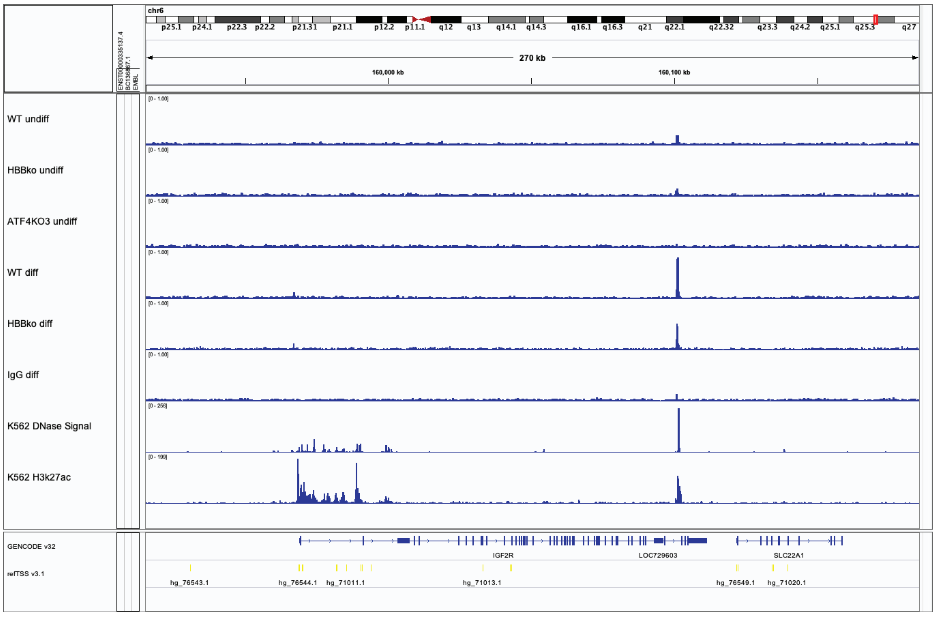

Supplement Fig 7

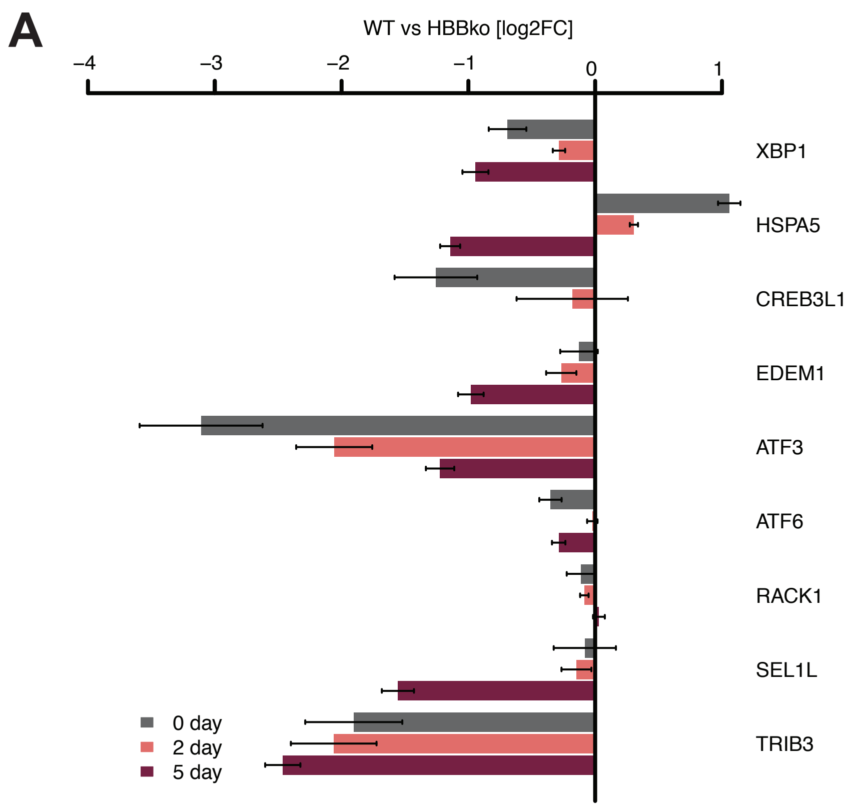
