## Supplemental Table 1 for "ATF4 mediates fetal globin upregulation in response to reduced β-globin"

| Identified Proteins (165/174) | Accession Number | Alternate ID | Molecular W | Exclusive Spectrum C peak_1 BioSample 1 | Exclusive Spectrum Cc peak_2 BioSample 2 | Exclusive Spectrum Cou peak_3 BioSample 3 | Exclusive Spectrum Co peak_4 BioSample 4 |
|---|---|---|---|---|---|---|---|
| Hemoglobin subunit alpha OS=Homo sapiens OX=9606 GN=HBA1 PE=1 SV=2 | HBA_HUMAN | HBA1 | 15 kDa | 31 | 88 | 108 | 224 |
| Cluster of Hemoglobin subunit beta OS=Homo sapiens OX=9606 GN=HBB PE=1 SV=2 (HBB_HUMAN) | HBB_HUMAN [2] | HBB | 16 kDa | 8 | 15 | 20 | 278 |
| Hemoglobin subunit beta OS=Homo sapiens OX=9606 GN=HBB PE=1 SV=2 | HBB_HUMAN | HBB | 16 kDa | 4 | 7 | 6 | 186 |
| Hemoglobin subunit delta OS=Homo sapiens OX=9606 GN=HBD PE=1 SV=2 | HBD_HUMAN | HBD | 16 kDa | 2 | 4 | 7 | 7 |
| Cluster of Hemoglobin subunit gamma-2 OS=Homo sapiens OX=9606 GN=HBG2 PE=1 SV=2 (HBG2_HUMAN) | HBG2_HUMAN [2] | HBG2 | 16 kDa | 43 | 146 | 79 | 10 |
| Hemoglobin subunit gamma-2 OS=Homo sapiens OX=9606 GN=HBG2 PE=1 SV=2 | HBG2_HUMAN | HBG2 | 16 kDa | 9 | 30 | 14 | 2 |
| Hemoglobin subunit gamma-1 OS=Homo sapiens OX=9606 GN=HBG1 PE=1 SV=2 | HBG1_HUMAN | HBG1 | 16 kDa | 8 | 18 | 10 | 1 |
| Glucose-6-phosphate isomerase OS=Homo sapiens OX=9606 GN=GPI PE=1 SV=4 | G6PI_HUMAN | GPI | 63 kDa | 0 | 0 | 12 | 39 |
| Aspartate aminotransferase, mitochondrial OS=Homo sapiens OX=9606 GN=GOT2 PE=1 SV=3 | AATM_HUMAN | GOT2 | 48 kDa | 0 | 10 | 23 | 2 |
| Tubulin alpha-1A chain OS=Homo sapiens OX=9606 GN=TUBA1A PE=1 SV=1 | TBA1A_HUMAN (+2) | TUBA1A | 50 kDa | 13 | 10 | 6 | 3 |
| Receptor of activated protein C kinase 1 OS=Homo sapiens OX=9606 GN=RACK1 PE=1 SV=3 | RACK1_HUMAN | RACK1 | 35 kDa | 3 | 2 | 14 | 12 |
| Cluster of Alpha-enolase OS=Homo sapiens OX=9606 GN=ENO1 PE=1 SV=2 (ENOA_HUMAN) | ENOA_HUMAN [2] | ENO1 | 47 kDa | 0 | 5 | 9 | 17 |
| Alpha-enolase OS=Homo sapiens OX=9606 GN=ENO1 PE=1 SV=2 | ENOA_HUMAN | ENO1 | 47 kDa | 0 | 4 | 7 | 15 |
| Gamma-enolase OS=Homo sapiens OX=9606 GN=ENO2 PE=1 SV=3 | ENOG_HUMAN | ENO2 | 47 kDa | 0 | 0 | 0 | 1 |
| GMP synthase [glutamine-hydrolyzing] OS=Homo sapiens OX=9606 GN=GMPS PE=1 SV=1 | GUAA_HUMAN | GMPS | 77 kDa | 0 | 16 | 12 | 0 |
| Peptidyl-prolyl cis-trans isomerase A OS=Homo sapiens OX=9606 GN=PPIA PE=1 SV=2 | PPIA_HUMAN | PPIA | 18 kDa | 14 | 12 | 2 | 0 |
| Carbonic anhydrase 2 OS=Homo sapiens OX=9606 GN=CA2 PE=1 SV=2 | CAH2_HUMAN | CA2 | 29 kDa | 13 | 5 | 4 | 5 |
| Phosphatidylethanolamine-binding protein 1 OS=Homo sapiens OX=9606 GN=PEBP1 PE=1 SV=3 | PEBP1_HUMAN | PEBP1 | 21 kDa | 19 | 5 | 3 | 0 |
| Fascin OS=Homo sapiens OX=9606 GN=FSCN1 PE=1 SV=3 | FSCN1_HUMAN | FSCN1 | 55 kDa | 0 | 2 | 12 | 12 |
| 60S ribosomal protein L7 OS=Homo sapiens OX=9606 GN=RPL7 PE=1 SV=1 | RL7_HUMAN | RPL7 | 29 kDa | 9 | 10 | 3 | 2 |
| Cluster of Tubulin beta chain OS=Homo sapiens OX=9606 GN=TUBB PE=1 SV=2 (TBB5_HUMAN) | TBB5_HUMAN [2] | TUBB | 50 kDa | 8 | 7 | 5 | 4 |
| Tubulin beta chain OS=Homo sapiens OX=9606 GN=TUBB PE=1 SV=2 | TBB5_HUMAN | TUBB | 50 kDa | 2 | 2 | 1 | 1 |
| Tubulin beta-4B chain OS=Homo sapiens OX=9606 GN=TUBB4B PE=1 SV=1 | TBB4B_HUMAN | TUBB4B | 50 kDa | 1 | 1 | 1 | 0 |
| Elongation factor 1-alpha 1 OS=Homo sapiens OX=9606 GN=EEF1A1 PE=1 SV=1 | EF1A1_HUMAN (+1) | EEF1A1 | 50 kDa | 9 | 6 | 6 | 2 |
| Heat shock cognate 71 kDa protein OS=Homo sapiens OX=9606 GN=HSPA8 PE=1 SV=1 | HSP7C_HUMAN | HSPA8 | 71 kDa | 7 | 6 | 6 | 3 |
| Proteasome subunit alpha type-7 OS=Homo sapiens OX=9606 GN=PSMA7 PE=1 SV=1 | PSA7_HUMAN | PSMA7 | 28 kDa | 4 | 7 | 4 | 7 |
| Malate dehydrogenase, cytoplasmic OS=Homo sapiens OX=9606 GN=MDH1 PE=1 SV=4 | MDHC_HUMAN | MDH1 | 36 kDa | 13 | 6 | 2 | 0 |
| Cofilin-1 OS=Homo sapiens OX=9606 GN=CFL1 PE=1 SV=3 | COF1_HUMAN | CFL1 | 19 kDa | 9 | 4 | 5 | 2 |
| Fumarate hydratase, mitochondrial OS=Homo sapiens OX=9606 GN=FH PE=1 SV=3 | FUMH_HUMAN | FH | 55 kDa | 7 | 10 | 2 | 0 |
| 60S ribosomal protein L6 OS=Homo sapiens OX=9606 GN=RPL6 PE=1 SV=3 | RL6_HUMAN | RPL6 | 33 kDa | 10 | 7 | 2 | 0 |
| Uroporphyrinogen decarboxylase OS=Homo sapiens OX=9606 GN=UROD PE=1 SV=2 | DCUP_HUMAN | UROD | 41 kDa | 0 | 5 | 13 | 0 |
| Histone H4 OS=Homo sapiens OX=9606 GN=HIST1H4A PE=1 SV=2 | H4_HUMAN | HIST1H4A | 11 kDa | 6 | 7 | 3 | 2 |
| Glycine--tRNA ligase OS=Homo sapiens OX=9606 GN=GARS PE=1 SV=3 | GARS_HUMAN | GARS | 83 kDa | 0 | 2 | 0 | 15 |
| Transketolase OS=Homo sapiens OX=9606 GN=TKT PE=1 SV=3 | TKT_HUMAN | TKT | 68 kDa | 0 | 0 | 0 | 17 |
| Histone H2A type 1-D OS=Homo sapiens OX=9606 GN=HIST1H2AD PE=1 SV=2 | H2A1D_HUMAN (+6) | HIST1H2AD | 14 kDa | 6 | 5 | 0 | 4 |
| Cluster of Histone H2B type 1-C/E/F/G/I OS=Homo sapiens OX=9606 GN=HIST1H2BC PE=1 SV=4 (H2B1C_HUMA | H2B1C_HUMAN [9] | HIST1H2BC | 14 kDa | 6 | 4 | 3 | 0 |
| Histone H2B type 1-C/E/F/G/I OS=Homo sapiens OX=9606 GN=HIST1H2BC PE=1 SV=4 | H2B1C_HUMAN (+8) | HIST1H2BC | 14 kDa | 3 | 2 | 1 | 0 |
| Proteasome subunit alpha type-4 OS=Homo sapiens OX=9606 GN=PSMA4 PE=1 SV=1 | PSA4_HUMAN | PSMA4 | 29 kDa | 2 | 5 | 4 | 5 |
| 60S ribosomal protein L7a OS=Homo sapiens OX=9606 GN=RPL7A PE=1 SV=2 | RL7A_HUMAN | RPL7A | 30 kDa | 8 | 6 | 2 | 0 |
| Cluster of Actin, cytoplasmic 1 OS=Homo sapiens OX=9606 GN=ACTB PE=1 SV=1 (ACTB_HUMAN) | ACTB_HUMAN [3] | ACTB | 42 kDa | 7 | 4 | 4 | 0 |
| Actin, cytoplasmic 1 OS=Homo sapiens OX=9606 GN=ACTB PE=1 SV=1 | ACTB_HUMAN (+1) | ACTB | 42 kDa | 5 | 2 | 3 | 0 |
| Beta-actin-like protein 2 OS=Homo sapiens OX=9606 GN=ACTBL2 PE=1 SV=2 | ACTBL_HUMAN | ACTBL2 | 42 kDa | 0 | 1 | 0 | 0 |
| Flavin reductase (NADPH) OS=Homo sapiens OX=9606 GN=BLVRB PE=1 SV=3 | BLVRB_HUMAN | BLVRB | 22 kDa | 0 | 3 | 4 | 8 |
| 60S ribosomal protein L4 OS=Homo sapiens OX=9606 GN=RPL4 PE=1 SV=5 | RL4_HUMAN | RPL4 | 48 kDa | 8 | 4 | 3 | 0 |
| 40S ribosomal protein S11 OS=Homo sapiens OX=9606 GN=RPS11 PE=1 SV=3 | RS11_HUMAN | RPS11 | 18 kDa | 6 | 5 | 4 | 0 |
| Oxygen-dependent coproporphyrinogen-III oxidase, mitochondrial OS=Homo sapiens OX=9606 GN=CPOX PE=1 S | HEM6_HUMAN | CPOX | 50 kDa | 0 | 4 | 6 | 3 |
| L-lactate dehydrogenase B chain OS=Homo sapiens OX=9606 GN=LDHB PE=1 SV=2 | LDHB_HUMAN | LDHB | 37 kDa | 7 | 3 | 4 | 0 |
| Protein disulfide-isomerase A3 OS=Homo sapiens OX=9606 GN=PDIA3 PE=1 SV=4 | PDIA3_HUMAN | PDIA3 | 57 kDa | 0 | 4 | 8 | 2 |
| Rab GDP dissociation inhibitor beta OS=Homo sapiens OX=9606 GN=GDI2 PE=1 SV=2 | GDIB_HUMAN | GDI2 | 51 kDa | 5 | 8 | 0 | 0 |
| Hydroxyacylglutathione hydrolase, mitochondrial OS=Homo sapiens OX=9606 GN=HAGH PE=1 SV=2 | GLO2_HUMAN | HAGH | 34 kDa | 0 | 0 | 3 | 10 |
| Phosphoglycerate kinase 1 OS=Homo sapiens OX=9606 GN=PGK1 PE=1 SV=3 | PGK1_HUMAN | PGK1 | 45 kDa | 4 | 0 | 8 | 0 |
| 60S ribosomal protein L18 OS=Homo sapiens OX=9606 GN=RPL18 PE=1 SV=2 | RL18_HUMAN | RPL18 | 22 kDa | 4 | 5 | 4 | 0 |
| 60S ribosomal protein L3 OS=Homo sapiens OX=9606 GN=RPL3 PE=1 SV=2 | RL3_HUMAN | RPL3 | 46 kDa | 6 | 5 | 2 | 0 |
| Porphobilinogen deaminase OS=Homo sapiens OX=9606 GN=HMBS PE=1 SV=2 | HEM3_HUMAN | HMBS | 39 kDa | 0 | 10 | 2 | 0 |
| L-lactate dehydrogenase A chain OS=Homo sapiens OX=9606 GN=LDHA PE=1 SV=2 | LDHA_HUMAN | LDHA | 37 kDa | 7 | 0 | 4 | 0 |
| 40S ribosomal protein S4, X isoform OS=Homo sapiens OX=9606 GN=RPS4X PE=1 SV=2 | RS4X_HUMAN | RPS4X | 30 kDa | 6 | 6 | 0 | 0 |
| 60S ribosomal protein L13 OS=Homo sapiens OX=9606 GN=RPL13 PE=1 SV=4 | RL13_HUMAN | RPL13 | 24 kDa | 5 | 4 | 2 | 0 |
| 40S ribosomal protein S8 OS=Homo sapiens OX=9606 GN=RPS8 PE=1 SV=2 | RS8_HUMAN | RPS8 | 24 kDa | 3 | 5 | 3 | 0 |
| Carbonic anhydrase 1 OS=Homo sapiens OX=9606 GN=CA1 PE=1 SV=2 | CAH1_HUMAN | CA1 | 29 kDa | 3 | 5 | 3 | 0 |
| Sorbitol dehydrogenase OS=Homo sapiens OX=9606 GN=SORD PE=1 SV=4 | DHSO_HUMAN | SORD | 38 kDa | 0 | 0 | 0 | 10 |
| Delta(3,5)-Delta(2,4)-dienoyl-CoA isomerase, mitochondrial OS=Homo sapiens OX=9606 GN=ECH1 PE=1 SV=2 | ECH1_HUMAN | ECH1 | 36 kDa | 2 | 3 | 4 | 0 |
| Histone H1.3 OS=Homo sapiens OX=9606 GN=HIST1H1D PE=1 SV=2 | H13_HUMAN | HIST1H1D | 22 kDa | 5 | 3 | 2 | 0 |
| UMP-CMP kinase OS=Homo sapiens OX=9606 GN=CMPK1 PE=1 SV=3 | KCY_HUMAN | CMPK1 | 22 kDa | 0 | 0 | 0 | 10 |
| Nucleolin OS=Homo sapiens OX=9606 GN=NCL PE=1 SV=3 | NUCL_HUMAN | NCL | 77 kDa | 10 | 0 | 0 | 0 |
| Proteasome subunit alpha type-1 OS=Homo sapiens OX=9606 GN=PSMA1 PE=1 SV=1 | PSA1_HUMAN | PSMA1 | 30 kDa | 3 | 5 | 0 | 2 |
| 40S ribosomal protein S16 OS=Homo sapiens OX=9606 GN=RPS16 PE=1 SV=2 | RS16_HUMAN | RPS16 | 16 kDa | 4 | 5 | 0 | 0 |
| 40S ribosomal protein S3 OS=Homo sapiens OX=9606 GN=RPS3 PE=1 SV=2 | RS3_HUMAN | RPS3 | 27 kDa | 5 | 3 | 2 | 0 |
| 60S ribosomal protein L15 OS=Homo sapiens OX=9606 GN=RPL15 PE=1 SV=2 | RL15_HUMAN | RPL15 | 24 kDa | 4 | 4 | 2 | 0 |
| Proteasome subunit alpha type-5 OS=Homo sapiens OX=9606 GN=PSMA5 PE=1 SV=3 | PSA5_HUMAN | PSMA5 | 26 kDa | 2 | 4 | 0 | 3 |
| Hepatoma-derived growth factor OS=Homo sapiens OX=9606 GN=HDGF PE=1 SV=1 | HDGF_HUMAN | HDGF | 27 kDa | 4 | 3 | 3 | 0 |
| 60S ribosomal protein L35a OS=Homo sapiens OX=9606 GN=RPL35A PE=1 SV=2 | RL35A_HUMAN | RPL35A | 13 kDa | 5 | 3 | 2 | 0 |
| 60S ribosomal protein L27 OS=Homo sapiens OX=9606 GN=RPL27 PE=1 SV=2 | RL27_HUMAN | RPL27 | 16 kDa | 5 | 5 | 0 | 0 |
| Histone H3.1t OS=Homo sapiens OX=9606 GN=HIST3H3 PE=1 SV=3 | H31T_HUMAN (+3) | HIST3H3 | 16 kDa | 4 | 3 | 3 | 0 |
| Heat shock 70 kDa protein 1A OS=Homo sapiens OX=9606 GN=HSPA1A PE=1 SV=1 | HS71A_HUMAN (+1) | HSPA1A | 70 kDa | 4 | 3 | 0 | 0 |
| 40S ribosomal protein S6 OS=Homo sapiens OX=9606 GN=RPS6 PE=1 SV=1 | RS6_HUMAN | RPS6 | 29 kDa | 4 | 3 | 2 | 0 |
| Peroxiredoxin-2 OS=Homo sapiens OX=9606 GN=PRDX2 PE=1 SV=5 | PRDX2_HUMAN | PRDX2 | 22 kDa | 2 | 1 | 4 | 2 |
| Ubiquitin-60S ribosomal protein L40 OS=Homo sapiens OX=9606 GN=UBA52 PE=1 SV=2 | RL40_HUMAN (+3) | UBA52 | 15 kDa | 2 | 2 | 3 | 2 |
| Adenylate kinase isoenzyme 1 OS=Homo sapiens OX=9606 GN=AK1 PE=1 SV=3 | KAD1_HUMAN | AK1 | 22 kDa | 0 | 0 | 0 | 8 |
| MOB kinase activator 1A OS=Homo sapiens OX=9606 GN=MOB1A PE=1 SV=4 | MOB1A_HUMAN (+1) | MOB1A | 25 kDa | 0 | 0 | 0 | 8 |
| Peptidyl-prolyl cis-trans isomerase NIMA-interacting 1 OS=Homo sapiens OX=9606 GN=PIN1 PE=1 SV=1 | PIN1_HUMAN | PIN1 | 18 kDa | 0 | 3 | 5 | 0 |
| Proteasome subunit beta type-5 OS=Homo sapiens OX=9606 GN=PSMB5 PE=1 SV=3 | PSB5_HUMAN | PSMB5 | 28 kDa | 0 | 5 | 0 | 2 |
| Proteasome subunit beta type-6 OS=Homo sapiens OX=9606 GN=PSMB6 PE=1 SV=4 | PSB6_HUMAN | PSMB6 | 25 kDa | 3 | 3 | 2 | 0 |
| Alanine--tRNA ligase, cytoplasmic OS=Homo sapiens OX=9606 GN=AARS PE=1 SV=2 | SYAC_HUMAN | AARS | 107 kDa | 0 | 8 | 0 | 0 |
| 60S ribosomal protein L13a OS=Homo sapiens OX=9606 GN=RPL13A PE=1 SV=2 | RL13A_HUMAN | RPL13A | 24 kDa | 3 | 3 | 2 | 0 |
| 60S ribosomal protein L8 OS=Homo sapiens OX=9606 GN=RPL8 PE=1 SV=2 | RL8_HUMAN | RPL8 | 28 kDa | 2 | 4 | 2 | 0 |
| Cluster of Ubiquitin-conjugating enzyme E2 variant 2 OS=Homo sapiens OX=9606 GN=UBE2V2 PE=1 SV=4 (UB2 | UB2V2_HUMAN | UBE2V2 | 16 kDa | 3 | 4 | 0 | 0 |
| Ubiquitin-conjugating enzyme E2 variant 2 OS=Homo sapiens OX=9606 GN=UBE2V2 PE=1 SV=4 | UB2V2_HUMAN | UBE2V2 | 16 kDa | 2 | 2 | 0 | 0 |
| Aconitate hydratase, mitochondrial OS=Homo sapiens OX=9606 GN=ACO2 PE=1 SV=2 | ACON_HUMAN | ACO2 | 85 kDa | 0 | 2 | 5 | 0 |
| Phosphoserine aminotransferase OS=Homo sapiens OX=9606 GN=PSAT1 PE=1 SV=2 | SERC_HUMAN | PSAT1 | 40 kDa | 2 | 5 | 0 | 0 |
| Fructose-bisphosphate aldolase A OS=Homo sapiens OX=9606 GN=ALDOA PE=1 SV=2 | ALDOA_HUMAN | ALDOA | 39 kDa | 0 | 0 | 0 | 4 |
| Proteasome subunit alpha type-6 OS=Homo sapiens OX=9606 GN=PSMA6 PE=1 SV=1 | PSA6_HUMAN | PSMA6 | 27 kDa | 2 | 3 | 0 | 0 |
| 40S ribosomal protein S9 OS=Homo sapiens OX=9606 GN=RPS9 PE=1 SV=3 | RS9_HUMAN | RPS9 | 23 kDa | 3 | 3 | 0 | 0 |
| Proteasome subunit beta type-2 OS=Homo sapiens OX=9606 GN=PSMB2 PE=1 SV=1 | PSB2_HUMAN | PSMB2 | 23 kDa | 0 | 3 | 2 | 2 |
| 40S ribosomal protein S14 OS=Homo sapiens OX=9606 GN=RPS14 PE=1 SV=3 | RS14_HUMAN | RPS14 | 16 kDa | 2 | 3 | 2 | 0 |
| 40S ribosomal protein S15a OS=Homo sapiens OX=9606 GN=RPS15A PE=1 SV=2 | RS15A_HUMAN | RPS15A | 15 kDa | 4 | 3 | 0 | 0 |
| Endoplasmic reticulum resident protein 29 OS=Homo sapiens OX=9606 GN=ERP29 PE=1 SV=4 | ERP29_HUMAN | ERP29 | 29 kDa | 0 | 6 | 0 | 0 |

| | | | | | | | |
|---|---|---|---|---|---|---|---|
| Tryptophan--tRNA ligase, cytoplasmic OS=Homo sapiens OX=9606 GN=WARS PE=1 SV=2 | SYWC_HUMAN | WARS | 53 kDa | 0 | 0 | 0 | 6 |
| Serotransferrin OS=Homo sapiens OX=9606 GN=TF PE=1 SV=3 | TRFE_HUMAN | TF | 77 kDa | 0 | 0 | 5 | 0 |
| 40S ribosomal protein S2 OS=Homo sapiens OX=9606 GN=RPS2 PE=1 SV=2 | RS2_HUMAN | RPS2 | 31 kDa | 4 | 0 | 0 | 0 |
| Heterogeneous nuclear ribonucleoprotein U OS=Homo sapiens OX=9606 GN=HNRNPU PE=1 SV=6 | HNRPU_HUMAN | HNRNPU | 91 kDa | 4 | 0 | 0 | 0 |
| Dermcidin OS=Homo sapiens OX=9606 GN=DCD PE=1 SV=2 | DCD_HUMAN | DCD | 11 kDa | 4 | 0 | 0 | 0 |
| 60S ribosomal protein L19 OS=Homo sapiens OX=9606 GN=RPL19 PE=1 SV=1 | RL19_HUMAN | RPL19 | 23 kDa | 3 | 2 | 0 | 0 |
| 60S ribosomal protein L26-like 1 OS=Homo sapiens OX=9606 GN=RPL26L1 PE=1 SV=1 | RL26L_HUMAN (+1) | RPL26L1 | 17 kDa | 4 | 2 | 0 | 0 |
| 60S ribosomal protein L27a OS=Homo sapiens OX=9606 GN=RPL27A PE=1 SV=2 | RL27A_HUMAN | RPL27A | 17 kDa | 3 | 0 | 0 | 0 |
| Cluster of Ubiquitin-conjugating enzyme E2 D3 OS=Homo sapiens OX=9606 GN=UBE2D3 PE=1 SV=1 (UB2D3_HU | UB2D3_HUMAN | UBE2D3 | 17 kDa | 0 | 3 | 2 | 0 |
| Ubiquitin-conjugating enzyme E2 D3 OS=Homo sapiens OX=9606 GN=UBE2D3 PE=1 SV=1 | UB2D3_HUMAN | UBE2D3 | 17 kDa | 0 | 2 | 1 | 0 |
| Poly(rC)-binding protein 1 OS=Homo sapiens OX=9606 GN=PCBP1 PE=1 SV=2 | PCBP1_HUMAN | PCBP1 | 37 kDa | 0 | 0 | 0 | 5 |
| Multifunctional protein ADE2 OS=Homo sapiens OX=9606 GN=PAICS PE=1 SV=3 | PUR6_HUMAN | PAICS | 47 kDa | 0 | 0 | 0 | 5 |
| Ribose-5-phosphate isomerase OS=Homo sapiens OX=9606 GN=RPIA PE=1 SV=3 | RPIA_HUMAN | RPIA | 33 kDa | 5 | 0 | 0 | 0 |
| Sialic acid synthase OS=Homo sapiens OX=9606 GN=NANS PE=1 SV=2 | SIAS_HUMAN | NANS | 40 kDa | 0 | 0 | 0 | 5 |
| Biliverdin reductase A OS=Homo sapiens OX=9606 GN=BLVRA PE=1 SV=2 | BIEA_HUMAN | BLVRA | 33 kDa | 0 | 0 | 2 | 3 |
| Selenide, water dikinase 1 OS=Homo sapiens OX=9606 GN=SEPHS1 PE=1 SV=2 | SPS1_HUMAN | SEPHS1 | 43 kDa | 0 | 0 | 0 | 5 |
| Ras suppressor protein 1 OS=Homo sapiens OX=9606 GN=RSU1 PE=1 SV=3 | RSU1_HUMAN | RSU1 | 32 kDa | 0 | 0 | 5 | 0 |
| 60S ribosomal protein L22 OS=Homo sapiens OX=9606 GN=RPL22 PE=1 SV=2 | RL22_HUMAN | RPL22 | 15 kDa | 3 | 0 | 0 | 0 |
| 10 kDa heat shock protein, mitochondrial OS=Homo sapiens OX=9606 GN=HSPE1 PE=1 SV=2 | CH10_HUMAN | HSPE1 | 11 kDa | 0 | 0 | 3 | 2 |
| Heterogeneous nuclear ribonucleoprotein A1 OS=Homo sapiens OX=9606 GN=HNRNPA1 PE=1 SV=5 | ROA1_HUMAN | HNRNPA1 | 39 kDa | 3 | 2 | 0 | 0 |
| Phosphoenolpyruvate carboxykinase [GTP], mitochondrial OS=Homo sapiens OX=9606 GN=PCK2 PE=1 SV=4 | PCKGM_HUMAN | PCK2 | 71 kDa | 0 | 0 | 0 | 5 |
| 60S ribosomal protein L21 OS=Homo sapiens OX=9606 GN=RPL21 PE=1 SV=2 | RL21_HUMAN | RPL21 | 19 kDa | 3 | 2 | 0 | 0 |
| GTP-binding nuclear protein Ran OS=Homo sapiens OX=9606 GN=RAN PE=1 SV=3 | RAN_HUMAN | RAN | 24 kDa | 3 | 2 | 0 | 0 |
| 40S ribosomal protein S26 OS=Homo sapiens OX=9606 GN=RPS26 PE=1 SV=3 | RS26_HUMAN | RPS26 | 13 kDa | 2 | 2 | 0 | 0 |
| 40S ribosomal protein S13 OS=Homo sapiens OX=9606 GN=RPS13 PE=1 SV=2 | RS13_HUMAN | RPS13 | 17 kDa | 3 | 0 | 0 | 0 |
| Heterogeneous nuclear ribonucleoproteins A2/B1 OS=Homo sapiens OX=9606 GN=HNRNPA2B1 PE=1 SV=2 | ROA2_HUMAN | HNRNPA2B1 | 37 kDa | 2 | 3 | 0 | 0 |
| Mth938 domain-containing protein OS=Homo sapiens OX=9606 GN=AAMDC PE=1 SV=1 | AAMDC_HUMAN | AAMDC | 13 kDa | 3 | 2 | 0 | 0 |
| 60S ribosomal protein L28 OS=Homo sapiens OX=9606 GN=RPL28 PE=1 SV=3 | RL28_HUMAN | RPL28 | 16 kDa | 4 | 0 | 0 | 0 |
| Nuclease-sensitive element-binding protein 1 OS=Homo sapiens OX=9606 GN=YBX1 PE=1 SV=3 | YBOX1_HUMAN | YBX1 | 36 kDa | 0 | 0 | 0 | 2 |
| Transgelin-2 OS=Homo sapiens OX=9606 GN=TAGLN2 PE=1 SV=3 | TAGL2_HUMAN | TAGLN2 | 22 kDa | 4 | 0 | 0 | 0 |
| SEC14-like protein 4 OS=Homo sapiens OX=9606 GN=SEC14L4 PE=1 SV=1 | S14L4_HUMAN | SEC14L4 | 47 kDa | 0 | 4 | 0 | 0 |
| Far upstream element-binding protein 1 OS=Homo sapiens OX=9606 GN=FUBP1 PE=1 SV=3 | FUBP1_HUMAN | FUBP1 | 68 kDa | 0 | 0 | 4 | 0 |
| Fumarylacetoacetase OS=Homo sapiens OX=9606 GN=FAH PE=1 SV=2 | FAAA_HUMAN | FAH | 46 kDa | 3 | 0 | 0 | 0 |
| Electron transfer flavoprotein subunit beta OS=Homo sapiens OX=9606 GN=ETFB PE=1 SV=3 | ETFB_HUMAN | ETFB | 28 kDa | 4 | 0 | 0 | 0 |
| 60S ribosomal protein L10 OS=Homo sapiens OX=9606 GN=RPL10 PE=1 SV=4 | RL10_HUMAN | RPL10 | 25 kDa | 4 | 0 | 0 | 0 |
| Sorting nexin-3 OS=Homo sapiens OX=9606 GN=SNX3 PE=1 SV=3 | SNX3_HUMAN | SNX3 | 19 kDa | 0 | 0 | 4 | 0 |
| Eukaryotic initiation factor 4A-I OS=Homo sapiens OX=9606 GN=EIF4A1 PE=1 SV=1 | IF4A1_HUMAN | EIF4A1 | 46 kDa | 4 | 0 | 0 | 0 |
| Elongation factor 2 OS=Homo sapiens OX=9606 GN=EEF2 PE=1 SV=4 | EF2_HUMAN | EEF2 | 95 kDa | 0 | 0 | 0 | 4 |
| Proteasome subunit beta type-4 OS=Homo sapiens OX=9606 GN=PSMB4 PE=1 SV=4 | PSB4_HUMAN | PSMB4 | 29 kDa | 0 | 0 | 0 | 3 |
| Jupiter microtubule associated homolog 2 OS=Homo sapiens OX=9606 GN=JPT2 PE=1 SV=1 | JUPI2_HUMAN | JPT2 | 20 kDa | 0 | 0 | 4 | 0 |
| Ubiquitin-conjugating enzyme E2 L3 OS=Homo sapiens OX=9606 GN=UBE2L3 PE=1 SV=1 | UB2L3_HUMAN | UBE2L3 | 18 kDa | 0 | 4 | 0 | 0 |
| Proteasome subunit alpha type-2 OS=Homo sapiens OX=9606 GN=PSMA2 PE=1 SV=2 | PSA2_HUMAN | PSMA2 | 26 kDa | 0 | 0 | 0 | 4 |
| Branched-chain-amino-acid aminotransferase, mitochondrial OS=Homo sapiens OX=9606 GN=BCAT2 PE=1 SV= | BCAT2_HUMAN | BCAT2 | 44 kDa | 4 | 0 | 0 | 0 |
| 40S ribosomal protein S25 OS=Homo sapiens OX=9606 GN=RPS25 PE=1 SV=1 | RS25_HUMAN | RPS25 | 14 kDa | 2 | 0 | 0 | 0 |
| 60S ribosomal protein L23 OS=Homo sapiens OX=9606 GN=RPL23 PE=1 SV=1 | RL23_HUMAN | RPL23 | 15 kDa | 2 | 2 | 0 | 0 |
| 60S ribosomal protein L31 OS=Homo sapiens OX=9606 GN=RPL31 PE=1 SV=1 | RL31_HUMAN | RPL31 | 14 kDa | 2 | 0 | 0 | 0 |
| Proteasome subunit alpha type-3 OS=Homo sapiens OX=9606 GN=PSMA3 PE=1 SV=2 | PSA3_HUMAN | PSMA3 | 28 kDa | 2 | 2 | 0 | 0 |
| ATP-dependent RNA helicase DDX39A OS=Homo sapiens OX=9606 GN=DDX39A PE=1 SV=2 | DX39A_HUMAN (+1) | DDX39A | 49 kDa | 0 | 0 | 2 | 0 |
| Malate dehydrogenase, mitochondrial OS=Homo sapiens OX=9606 GN=MDH2 PE=1 SV=3 | MDHM_HUMAN | MDH2 | 36 kDa | 0 | 2 | 2 | 0 |
| Polyadenylate-binding protein 1 OS=Homo sapiens OX=9606 GN=PABPC1 PE=1 SV=2 | PABP1_HUMAN | PABPC1 | 71 kDa | 4 | 0 | 0 | 0 |
| Coiled-coil domain-containing protein 58 OS=Homo sapiens OX=9606 GN=CCDC58 PE=1 SV=1 | CCD58_HUMAN | CCDC58 | 17 kDa | 0 | 0 | 3 | 0 |
| 60S ribosomal protein L24 OS=Homo sapiens OX=9606 GN=RPL24 PE=1 SV=1 | RL24_HUMAN | RPL24 | 18 kDa | 3 | 0 | 0 | 0 |
| 60S ribosomal protein L23a OS=Homo sapiens OX=9606 GN=RPL23A PE=1 SV=1 | RL23A_HUMAN | RPL23A | 18 kDa | 3 | 0 | 0 | 0 |
| Dihydropteridine reductase OS=Homo sapiens OX=9606 GN=QDPR PE=1 SV=2 | DHPR_HUMAN | QDPR | 26 kDa | 0 | 0 | 3 | 0 |
| 60S ribosomal protein L34 OS=Homo sapiens OX=9606 GN=RPL34 PE=1 SV=3 | RL34_HUMAN | RPL34 | 13 kDa | 2 | 0 | 0 | 0 |
| DCN1-like protein 1 OS=Homo sapiens OX=9606 GN=DCUN1D1 PE=1 SV=1 | DCNL1_HUMAN | DCUN1D1 | 30 kDa | 0 | 0 | 3 | 0 |
| Heterogeneous nuclear ribonucleoprotein D0 OS=Homo sapiens OX=9606 GN=HNRNPD PE=1 SV=1 | HNRPD_HUMAN | HNRNPD | 38 kDa | 2 | 0 | 0 | 0 |
| 60S ribosomal protein L11 OS=Homo sapiens OX=9606 GN=RPL11 PE=1 SV=2 | RL11_HUMAN | RPL11 | 20 kDa | 2 | 0 | 0 | 0 |
| Dihydrolipoyl dehydrogenase, mitochondrial OS=Homo sapiens OX=9606 GN=DLD PE=1 SV=2 | DLDH_HUMAN | DLD | 54 kDa | 0 | 0 | 0 | 3 |
| 60S ribosomal protein L10a OS=Homo sapiens OX=9606 GN=RPL10A PE=1 SV=2 | RL10A_HUMAN | RPL10A | 25 kDa | 2 | 0 | 0 | 0 |
| Serum albumin OS=Homo sapiens OX=9606 GN=ALB PE=1 SV=2 | ALBU_HUMAN | ALB | 69 kDa | 2 | 0 | 0 | 0 |
| 26S proteasome non-ATPase regulatory subunit 9 OS=Homo sapiens OX=9606 GN=PSMD9 PE=1 SV=3 | PSMD9_HUMAN | PSMD9 | 25 kDa | 0 | 0 | 0 | 3 |
| Glutathione peroxidase 1 OS=Homo sapiens OX=9606 GN=GPX1 PE=1 SV=4 | GPX1_HUMAN | GPX1 | 22 kDa | 0 | 0 | 3 | 0 |
| Bifunctional purine biosynthesis protein PURH OS=Homo sapiens OX=9606 GN=ATIC PE=1 SV=3 | PUR9_HUMAN | ATIC | 65 kDa | 0 | 0 | 0 | 2 |
| Ubiquitin-conjugating enzyme E2 N OS=Homo sapiens OX=9606 GN=UBE2N PE=1 SV=1 | UBE2N_HUMAN | UBE2N | 17 kDa | 0 | 2 | 0 | 0 |
| Catalase OS=Homo sapiens OX=9606 GN=CAT PE=1 SV=3 | CATA_HUMAN | CAT | 60 kDa | 0 | 0 | 2 | 0 |
| 40S ribosomal protein S12 OS=Homo sapiens OX=9606 GN=RPS12 PE=1 SV=3 | RS12_HUMAN | RPS12 | 15 kDa | 0 | 0 | 0 | 2 |
| Polypyrimidine tract-binding protein 1 OS=Homo sapiens OX=9606 GN=PTBP1 PE=1 SV=1 | PTBP1_HUMAN | PTBP1 | 57 kDa | 0 | 0 | 2 | 0 |
| 40S ribosomal protein S18 OS=Homo sapiens OX=9606 GN=RPS18 PE=1 SV=3 | RS18_HUMAN | RPS18 | 18 kDa | 2 | 0 | 0 | 0 |
| Glyoxylate reductase/hydroxypyruvate reductase OS=Homo sapiens OX=9606 GN=GRHPR PE=1 SV=1 | GRHPR_HUMAN | GRHPR | 36 kDa | 0 | 0 | 0 | 2 |
| NEDD8-conjugating enzyme Ubc12 OS=Homo sapiens OX=9606 GN=UBE2M PE=1 SV=1 | UBC12_HUMAN | UBE2M | 21 kDa | 0 | 2 | 0 | 0 |
| Glyceraldehyde-3-phosphate dehydrogenase OS=Homo sapiens OX=9606 GN=GAPDH PE=1 SV=3 | G3P_HUMAN | GAPDH | 36 kDa | 2 | 0 | 0 | 0 |
| Galectin-3 OS=Homo sapiens OX=9606 GN=LGALS3 PE=1 SV=5 | LEG3_HUMAN | LGALS3 | 26 kDa | 0 | 0 | 2 | 0 |
| Probable ATP-dependent RNA helicase DDX46 OS=Homo sapiens OX=9606 GN=DDX46 PE=1 SV=2 | DDX46_HUMAN | DDX46 | 117 kDa | 0 | 0 | 2 | 0 |
| Transcription intermediary factor 1-beta OS=Homo sapiens OX=9606 GN=TRIM28 PE=1 SV=5 | TIF1B_HUMAN | TRIM28 | 89 kDa | 2 | 0 | 0 | 0 |
| Peroxiredoxin-1 OS=Homo sapiens OX=9606 GN=PRDX1 PE=1 SV=1 | PRDX1_HUMAN | PRDX1 | 22 kDa | 0 | 0 | 1 | 0 |
| 60S ribosomal protein L30 OS=Homo sapiens OX=9606 GN=RPL30 PE=1 SV=2 | RL30_HUMAN | RPL30 | 13 kDa | 1 | 0 | 0 | 0 |
