## Supplemental Table 2 for "ATF4 mediates fetal globin upregulation in response to reduced β-globin"

| IVT assembly of sgRNA | sequence |
|---|---|
| e66 (g5) | GGATCCTAATACGACTCACTATAG**CATGGTGCATCTGACTCCTG**GTTTTAGAGCTAGAA |
| e87 (g10) | GGATCCTAATACGACTCACTATAG**CTTGCCCCACAGGGCAGTAA**GTTTTAGAGCTAGAA |
| i161 | GGATCCTAATACGACTCACTATAG**TGGTATCAAGGTTACAAGAC**GTTTTAGAGCTAGAA |
| i251 | GGATCCTAATACGACTCACTATAG**GGGTGGGAAAATAGACCAAT**GTTTTAGAGCTAGAA |
| e297 | GGATCCTAATACGACTCACTATAG**TGGTCTACCCTTGGACCCAG**GTTTTAGAGCTAGAA |
| e334 | GGATCCTAATACGACTCACTATAG**GCCCATAACAGCATCAGGAG**GTTTTAGAGCTAGAA |
| e383 | GGATCCTAATACGACTCACTATAG**GCTCATGGCAAGAAAGTGCT**GTTTTAGAGCTAGAA |
| e439 | GGATCCTAATACGACTCACTATAG**CAGCTCACTCAGTGTGGCAA**GTTTTAGAGCTAGAA |
| e1353 | GGATCCTAATACGACTCACTATAG**CACAGACCAGCACGTTGCCC**GTTTTAGAGCTAGAA |
| e1427 | GGATCCTAATACGACTCACTATAG**GCAGGCTGCCTATCAGAAAG**GTTTTAGAGCTAGAA |
| ATF4dN g1 | GGATCCTAATACGACTCACTATAG**AGTCCCGCCTCATAAGTGGA**GTTTTAGAGCTAGAA |
| ATF4dN g2 | GGATCCTAATACGACTCACTATAG**TAGATCCCACCAGGACGATG**GTTTTAGAGCTAGAA |
| ATF4 sgRNA 2 | GGATCCTAATACGACTCACTATAG**CTCGTCACAGCTACGCCCT**GTTTTAGAGCTAGAA |
| ATF4 sgRNA 3 | GGATCCTAATACGACTCACTATAG**TGGCCAACTATACGGCTCCA**GTTTTAGAGCTAGAA |

| **CRISPRi guides** | |
|---|---|
| HBB g1 | TAGACCACCAGCAGCCTAA |
| HBB g2 | GAACTTCAGGGTGAGTCTA |
| Non-targeting 1 | CGCCAAACGTGCCCTGACGG |
| Non-targeting 2 | GCTCGGTCCCGCGTCGTCG |
| SMG6 | ctcctccccgctcggccct |
| XRN1 | CTCGAAAGCCCCAGCTCTA |
| CD55 | GCTGCGACTCGGCGGAGTCC |
| CD59 | GCGCAGAAGCGGCTCGAGGC |

| **qRT-PCR primers** | |
|---|---|
| HBB qpcr for | TGTCCACTCCTGATGCTGTTATG |
| HBB qpcr rev | GGCACCGAGCACTTTCTTG |
| HBG1/2 qpcr for | CCTGTCCTCTGCCTCTGCC |
| HBG1/2 qpcr rev | GGATTGCCAAAACGGTCAC |
| HBA qpcr for | GGGTGGACCCGGTCAACTT |
| HBA qpcr rev | GAGGTGGGCGGCCAGGGT |
| HBE qpcr for | TGCTGAGGAGAAGGCTGCCG |
| HBE qpcr rev | TGGGTCCAGGGGTAAACAACGAGG |
| HBZ qpcr for | GAGGACCATCATTGTGTCCA |

| | |
|---|---|
| HBZ qpcr rev | AGTGCGGGAAGTAGGTCTTG |
| GAPDH qpcr for | CAACAGCGACACCCACTCCT |
| GAPDH qpcr rev | CACCCTGTTGCTGTAGCCAAA |
| SMG6 qpcr for | GCACGCATGGTGAACAGAAA |
| SMG6 qpcr rev | TGTTCAGCTCTCATTCTCGTCC |
| XRN1 qpcr for | TGGCTTACTGGTACATGGGC |
| XRN1 qpcr rev | TCATGCCTAAGGAGCCCAAC |
| ASNS qpcr for | ACTGGCTGCTAGAAAGGTGG |
| ASNS qpcr rev | GCCTGAATGCCTTCCTCAGA |

**CHIP-qPCR primers**

| | |
|---|---|
| ASNS promoter for | ATGATGAAACTTCCCGCACG |
| ASNS promoter rev | GGGATGTGGACAGCTTGACG |

**PCR primers for knockout validation**

| | |
|---|---|
| ATF4 WT for | CATTCCTCGATTCCAGCAAAGC |
| ATF4 WT rev | TGAGTGATGGGGCCAAGTGAG |
| ATF4 knockout for | CGTCCTCGGCCTTCACAATA |
| ATF4 knockout rev | TCTTCAGGATGAGGCTTCTGC |
| ATF4dN for | CCT GGT CTC CGT GAG CGT C |
| ATF4dN rev | CATCCAATCTGTCCCGGAGAAGG |

**Primers for TIDE Amplification and Analysis**

| | |
|---|---|
| HBB exon 1-2 for | ATG CTT AGA ACC GAG GTA GAG TTT |
| HBB exon 1-2 rev | CCT GAG ACT TCC ACA CTG ATG |
| HBB exon 3 for | AGGCAGAATCCAGATGCTCAAGGC |
| HBB exon 3 rev | GCA CGT GGA TCC TGA GAA CTT CAG |
